# Adaptive evolution at mRNA editing sites in soft-bodied cephalopods

**DOI:** 10.1101/2020.08.16.252932

**Authors:** Mikhail A. Moldovan, Zoe S. Chervontseva, Georgii A. Bazykin, Mikhail S. Gelfand

## Abstract

The bulk of variability in mRNA sequence arises due to mutation – change in DNA sequence which is heritable if it occurs in the germline. However, variation in mRNA can also be achieved by post-translational modification including mRNA editing, changes in mRNA nucleotide sequence that mimic the effect of mutations. Such modifications are not inherited directly; however, as the processes affecting them are encoded in the genome, they have a heritable component, and therefore can be shaped by selection. In soft-bodied cephalopods, adenine-to-inosine RNA editing is very frequent, and much of it occurs at nonsynonymous sites, affecting the sequence of the encoded protein. Here, we show that mRNA editing of individual nonsynonymous sites in cephalopods originates in evolution through substitutions at regions adjacent to these sites. As such substitutions mimic the effect of the substitution at the edited site itself, we hypothesize that they are favored by selection if the inosine is selectively advantageous to adenine at the edited position. Consistent with this hypothesis and with observations on *Drosophila* and human editing sites, we show that edited adenines are more frequently substituted with guanine, an informational analogue of inosine, in the course of evolution than their unedited counterparts, and for heavily edited adenines, these transitions are favored by positive selection. Thus, our study shows that coleoid editing sites may enhance adaptation, which, together with distinct evolutionary features of *Drosophila* and human editing sites, points at a general role of RNA editing in the molecular evolution of metazoans.

## Introduction

The process of natural selection requires heritable variation to be present in a population and the absence of genetic variants selection could act upon is generally considered to be a factor hampering adaptation (Lush 1937; Smith 1976; Barton and Partridge 2000; Lanfear et al. 2014; Rousselle et al. 2020). Heritable variation is generated mainly by the mutational process (Lewontin 1964; Avery and Hill 1977; Lynch and Walsh 1998). Hence, the mutation rate may be a factor affecting the evolution rate, which we, following J. Maynard Smith, define here as the rate of accumulation of beneficial mutations (Smith 1976; Nam et al. 2017; Rousselle et al. 2020). As shown recently, in populations with low genetic variability the mutation rate is indeed correlated with the evolution rate (Rousselle et al. 2020). Thus, in order to adapt, a low-polymorphic population may need additional expressed genetic variability. Here, we test the hypothesis that a potential source of such variability could be introduced by heritable epigenetic modifications, specifically, mRNA editing (Bass and Weintraub 1988; Gommans et al. 2009; Klironomos et al. 2013; Kronholm and Collins 2015).

We consider the A-to-I mRNA editing, where adenine (A) is modified to inosine (I) that is subsequently read by the translation machinery as guanine (G) (Bass and Weintraub 1988). In most of the studied organisms, the A-to-I editing affecting protein sequences is restricted to only a few thousand adenines, with the vast majority of edited adenines located in non-coding regions, e.g. in Alu-repeats (Kim 2004; Ramaswami et al. 2012; Yablonovitch et al. 2017). Edited sites are poorly conserved between species, suggesting that most editing events are non-functional, with a few possible exceptions (Yang et al. 2008; Pinto et al. 2014; Yu et al. 2016). However, in coleoids, soft-bodied cephalopods, about 1% of adenines in the transcriptome are edited, and re-coding (i.e., affecting the amino acid sequence) and conserved sites comprise considerable fractions (Alon et al. 2015; Liscovitch-Brauer et al. 2017). One explanation for this phenomenon comes from the observation that the conserved editing sites tend to be edited in the nervous tissue, and editing may contribute to the increased plasticity and complexity of the coleoid nervous system and behavior compared to other extant cephalopods (*Nautilus*) (Albertin et al. 2015; Alon et al. 2015; Liscovitch-Brauer et al. 2017; Eisenberg and Levanon 2018). This hypothesis is supported by analogous observations in other organisms (Pinto et al. 2014; Yu et al. 2016) and, although indirectly, by the finding that the A-to-I RNA editing has emerged approximately at the same time as the nervous systems of multicellular organisms have become more complex (Jin et al. 2009).

A-to-I editing is not absolutely efficient and, if it occurs at a non-synonymous site, would result in two non-identical proteins with a varying ratio (Gommans et al. 2009; Liscovitch-Brauer et al. 2017; Yablonovich et al. 2017). The efficiency of mRNA editing depends on the strength of the site motif and the local mRNA secondary structure (Morse et al. 2002; Reenan 2005; Gommans et al. 2009; Alon et al. 2012; Savva et al. 2012; Klironomos et al. 2013; Rieder et al. 2013; Liscovitch-Brauer et al. 2017). As the sequence and structure requirements seem to be relatively weak, mRNA editing sites have been proposed to constantly emerge at random points of the genome (Gommans et al. 2009; Xu and Zhang 2014).

To date, four models of A-to-I editing site evolution have been proposed. (i) Most A-to-I editing sites generally are not adaptive and mainly arise at positions with tolerable, i.e. effectively neutral or mildly deleterious, A-to-G substitutions (Xu and Zhang 2014). (ii) A-to-I editing is a mechanism of rescuing deleterious G-to-A substitutions (Jiang and Zhang 2019). (iii) A-to-I editing, generating multiple protein variants, is important for the advantageous transcriptome diversification, and hence the individual sites should be conserved (Liscovitch-Brauer et al. 2017; Eisenberg and Levanon 2018). (iv) The potential of A-to-I editing to mimic A-to-G substitutions is advantageous, and thus A-to-I editing sites function as transitory states when an advantageous mutation has not yet occurred (Popitsch et al., 2020).

Editing site evolution in *Drosophila* and human has been recently shown to adhere to model (iv) (Popitsch et al., 2020), while editing sites in coleoids are largely considered as means for proteome diversification as in model (iii) (Liscovitch-Brauer et al. 2017; Eisenberg and Levanon 2018) or be selectively neutral (Jiang and Zhang 2019). We attempt to resolve this controversy by detailed analysis of substitution patterns and selection regimes, taking into account the varying strength of A-to-I editing at different sites.

Generally, in a population with low genetic variability, one might expect evolutionary benefits of A-to-I editing consistent with model (iv). Indeed, if there is a position in the genome occupied by an adenine, but guanine in this position would yield a fitter genotype, there are two evolutionary pathways for adaptation: through an A-to-G substitution at this site, or through emergence of a local sequence context yielding or reinforcing A-to-I editing of this site. If the selective benefit conferred by both pathways is comparable, which of them will be taken will depend on the probability of the corresponding mutation (Yampolsky and Stolzfus 2001). A specific mutation is needed in the first scenario; by contrast, many different editing context-improving mutations could yield a fitter genotype, and the waiting time for any such mutation could be shorter (Durrett and Schmidt 2008). As a result, selection would lead to emergence of the adaptive editing phenotype.

We propose that non-conserved coleoid A-to-I mRNA editing sites, comprising the larger percentage relative to the conserved ones, could function as substitutes of beneficial A-to-G substitutions in low-polymorphic coleoid populations. We show that the levels of cephalopod A-to-I editing heavily depend on the sequence of adjacent regions, and hence are influenced by a multitude of possible mutations. Critically, we show that edited adenines are more frequently substituted in related species to guanines and less frequently, to cytosines or thymines, than non-edited ones. At strongly edited sites, the adenine-to-guanine transitions are favored by positive selection. Our results suggest that, while conserved coleoid editing sites could be functionally important *per se*, a large subset of non-conserved editing sites could play a role in the adaptive evolution by introducing, at least in a fraction of transcripts, guanines that are beneficial at the given positions. When this study had been completed, a similar observation was made for *Drosophila* and human editing sites by analysis of genomic polymorphisms (Popitsch et al., 2020). This indicates that A-to-I editing could have similar, imporant evolutionary roles in multiple metazoan lineages.

## Results

### Editing level is associated with the local and global sequence context

We studied the A-to-I editing using available genomic read libraries, transcriptomes, and editing sites data for four coleoids, closely related octopuses *Octopus vulgaris* and *O. bimaculoides*, squid *Loligo pealei*, and cuttlefish *Sepia esculenta* (Liscovitch-Brauer et al. 2017). As outgroups, we considered nautiloid *Nautilus pompilius* and gastropod mollusk *Aplysia californica* (Liscovitch-Brauer et al. 2017) (Fig. 1a).

**Figure 1.**
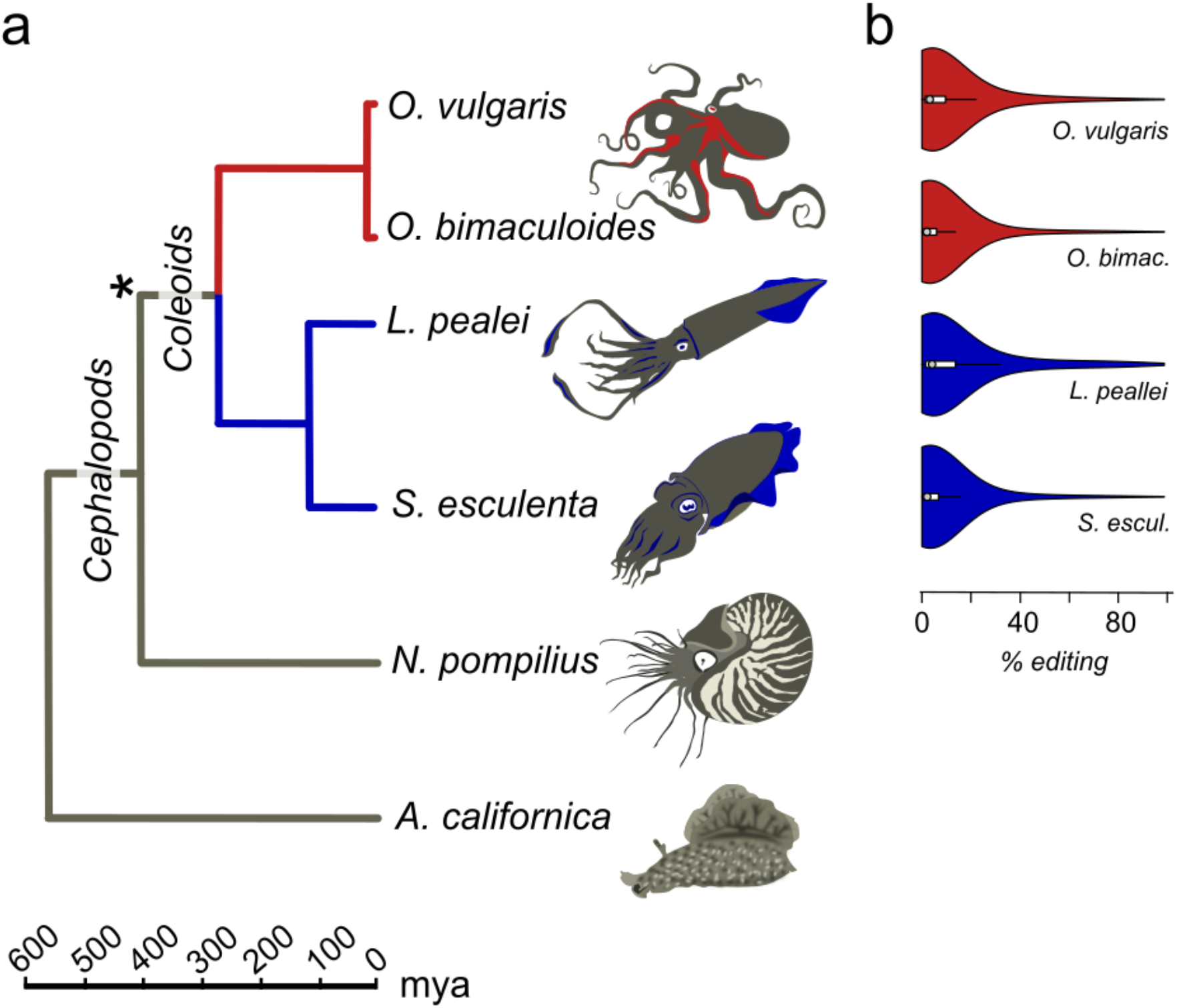
Prevalent mRNA editing in coleoid molluscs. (**a**) Phylogenetic tree of the species taken from TimeTree (Hedges et al. 2006). The asterisk marks the putative beginning of editing site expansion (Liscovitch-Brauer et al. 2017). (**b**) Distributions of per-nucleotide editing levels of the predicted editing sites in the studied coleoids.

The action of editing sites as surrogates of beneficial A-to-G substitutions presumes advantageous enhancement of editing probabilities at individual sites. As A-to-I editing is to be affected by the local sequence context (Alon et al. 2012; Liscovitch-Brauer et al. 2017) and the RNA secondary RNA structure (Morse et al. 2002; Reenan 2005; Gommans et al. 2009; Savva et al. 2012; Klironomos et al. 2013; Rieder et al. 2013), one would expect, firstly, contextual differences around weakly *vs*. heavily edited sites and, secondly, contextual mutations associated with changes in editing status. Indeed, we have observed a previously unnoted dependence of the editing level (EL) (Fig. 1b), defined as the percent of transcripts containing I at the considered site at the moment of sequencing (Fig. 2a, Suppl. Fig. S1), on the site context (±1 motif). Certain changes in the ±1 motif, specifically, an increase in the preference for G or T at the +1 position, are associated with the increase of EL, although its information content of the motif remains approximately the same. Although the ±1 motif of both weakly and strongly edited sites is consistent with the ADAR (adenosine deaminases acting on RNA) profile (Alon et al. 2012; Liscovitch-Brauer et al. 2017), this observation could point to the action of different ADAR enzymes or to different modes of action of the same enzyme on strongly and weakly edited sites. There also seem to be some differences between the motifs of conserved and non-conserved sites (Suppl. Fig. S2).

**Figure 2.**
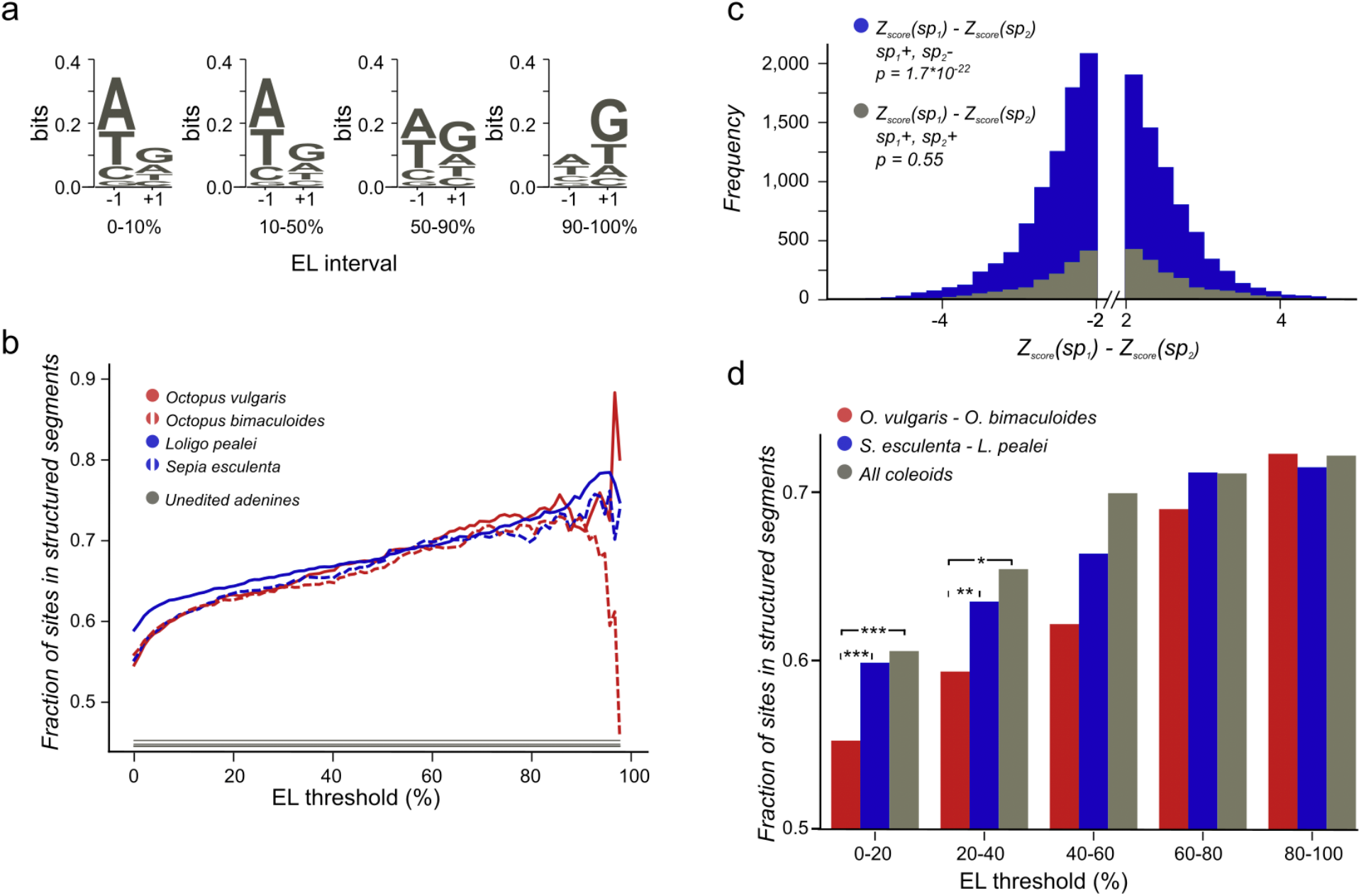
(**a**) *O. bimaculoides* editing site context changes with the increase of editing level. The height of the letters represents the LOGO bit score of each nucleotide. (**b**) Highly conserved editing sites tend to be relatively more structured. The fraction of editing sites that are in structured segments is shown for different editing levels: red — *O. vulgaris*, red dashed — *O. bimaculoides*, blue — *L. pealei*, blue dashed — *S. esculenta*, grey — the constant showing the fraction of unedited adenines located in structured segments. The noisy pattern at the right is due to a low number of very highly edited sites. (**c**) The stability of the local secondary structure is higher at edited adenines than at homologous, non-edited adenines for the squid/cuttlefish pair. The distribution of the difference of the minimal free energy Z-score between homologous sites in squid and cuttlefish is shown in blue when two homologous sites have different editing status (edited minus unedited) and in grey when both sites in a pair are edited. The left tail of the blue histogram is heavier than the right one (*p* = 9.37×10^−33^ versus 0.32 for the grey histogram), showing that the editing sites tend to regions with higher secondary structure stability. (**d**). Conserved editing sites tend to be more structured. The three groups of sites are those present in two of the four species *(O. vulgaris* and *O. bimaculoides*, red, or *L. pealei* and *S. esculenta,* blue), or in all four species (grey). Statistically significant differences are shown with brackets (*** *p*<0.001, * *p*<0.05).

The analysis of non-conserved editing sites (NCES) in the octopus pair demonstrates overrepresentation of mismatches in the ±1 motif of the edited adenines reinforcing the local editing context compared to the homologous unedited adenines, for which the editing context is not observed (See Materials and Methods, Suppl. Fig. S3). Thus, both the editing status and the EL of a site are associated with substitutions in the ±1 motif. In the squid-cuttlefish pair, the higher number of mutations obscures this analysis.

To estimate the size of the region that affects editing, we have measured the correlation between the editing level difference in conserved editing sites (CES) in closely related species and the number of mismatches in variable-sized windows centered at edited adenines. The window size yielding the largest correlation coefficient shows the average span of the context affecting the ADAR performance. For the *Octopus* pair, the highest correlation has been obtained at the window size of ~100 nucleotides (Suppl. Fig. S4), consistent with previous estimates for the length of the region affecting editing (Liscovitch-Brauer et al. 2017).

### Editing level is affected by secondary structure in adjacent RNA

The A-to-I editing in model species depends on large RNA structures spanning hundreds of nucleotides in addition to the local sequence context (Morse et al. 2002; Reenan 2005; Ensterö et al. 2009; Rieder et al. 2013; Kurmangaliyev et al. 2015) as the ADAR-mediated mRNA editing generally requires secondary RNA structures (Gommans et al. 2009; Farajollahi and Maas 2010; Xu and Zhang 2014). Thus, we have assessed the link between RNA secondary structure and ELs of focal sites.

We have predicted structured segments in the transcripts of all six considered species. As the fraction of adenines located within structured segments is the same for all cephalopod species, including *Nautilus* (Suppl. Fig. S5), our secondary structure analyses are not systematically influenced by the GC-content of the studied genomes (Wang et al. 1984). Then we have assessed the contribution of mRNA secondary structure to the editing process by comparing structural contexts of edited and unedited adenines (Materials and Methods). The fraction of edited adenines located in putative structured regions is higher than the respective fraction for non-edited sites. Moreover, sites that are more highly edited (Fig. 2b) as well as sites conserved between more distant species (Fig. 2d) tend to be more structured.

To uncover the connection between the strength of a local secondary RNA structure and the editing status at individual sites, we have compared the fractions of non-conserved editing sites (NCES) located within structured segments in edited *vs*. non-edited states. We considered the *Octopus* pair and the squid–cuttlefish pair. For both pairs, we have compared CES and NCES. For NCES in both species pairs we have observed significantly more cases when the edited site in a pair is more structured than the unedited one while the control CES set shows no bias (binomial test *p*<10^−3^ for all pairs; Fig. 2c, Suppl. Fig. S6).

Not only the fact of editing, but the difference in editing levels is linked to local secondary structures. For the closely related *Octopus* pair, we have calculated correlations between differences in ELs of homologous edited adenines and differences in their structure Z-scores (Suppl. Fig. S7). Almost no correlation (*r*=0.1, t-test *p* < 0.05) is seen when the EL difference is small (>5%), whereas for large differences in ELs (>50%) the correlation is substantial (*r*=0.7, t-test *p* < 0.05). A likely explanation is that larger differences are indeed due to the strength of the local secondary structure, whereas small differences in ELs arise as consequences of random noise. Consistent with the observations above, if we consider structures around edited adenines and their unedited homologs, setting the ELs of unedited adenines to 0, we observe a similar, although a weaker trend (Suppl. Fig. S7), with correlations reaching 0.4 (t-test *p* < 0.05) when the ELs of NCES are high.

The observations about local contexts, both the ±1 motif and RNA structures, imply that mutations near editing sites influence the editing status as well as the editing level.

### Edited adenines are often substituted by guanines

If edited adenines indeed frequently mimic the beneficial guanine state, the substitution patterns of edited and unedited adenines should differ, with edited adenines being more prone to substitutions to guanine and less prone to substitutions to cytosine or thymine (Popitsch et al. 2020). Firstly, we performed the analysis of the species pairs to infer the properties of A-G mismatches at editing sites. For a pair of considered species, we define *R* as the mismatch probability for an edited adenine divided by the probability of the same mismatch for an unedited adenine: *R_N_*=*p*(E,*N*)/*p*(A,*N*), where E and A are, respectively, edited and not edited adenines in one species, and *N* is the non-E, non-A nucleotide at the homologous site in the other organism. Similar formulas are applied when we consider directed substitutions instead of mismatches. If a pair of the ancestral and the descendant species is considered, we use notation *R*_→*N*_ to identify the directionality. *R*_→*N*_ = *p*(E→*N*)/*p*(A→*N*), where *p*(E→*N*) and *p*(A→*N*) are, respectively, the probabilities of the substitution of the edited and non-edited adenine to *N*. Similarly, notation *R*_*N*→_ is used when substitutions from ancestral *N* to E and A are considered: *R*_*N*→_=*p*(*N*→E)/*p*(*N*→A). Higher values of *R*_*N*→_ imply that the ancestral nucleotide *N* is more likely to be substituted by an edited adenine, compared to an unedited one.

We have observed a striking dependence of the calculated mismatch probabilities on the editing status of the adenines and their ELs. In the *Octopus* pair, *R*_G_ and *R*_Y_ (Y denotes pyrimidine, C or T) differ both in value and in the dependence on the EL (Fig. 3ab). Indeed, *R*_G_ is always higher than *R*_Y_ with *R*_G_ further increasing and *R*_Y_ decreasing as the EL increases. The probability for an adenine to be substituted by a guanine in the *O. vulgaris* lineage is ~8 times higher when the homologous adenine is strongly edited in *O. bimaculoides* than when it is not (Fig. 3a). For the more distantly related squid–cuttlefish pair, we observe a similar although less pronounced effect. For all distant pairs, that is, *Octopus*–squid/cuttlefish, *R*_G_ shows no or only a weak dependence on the EL.

**Figure 3.**
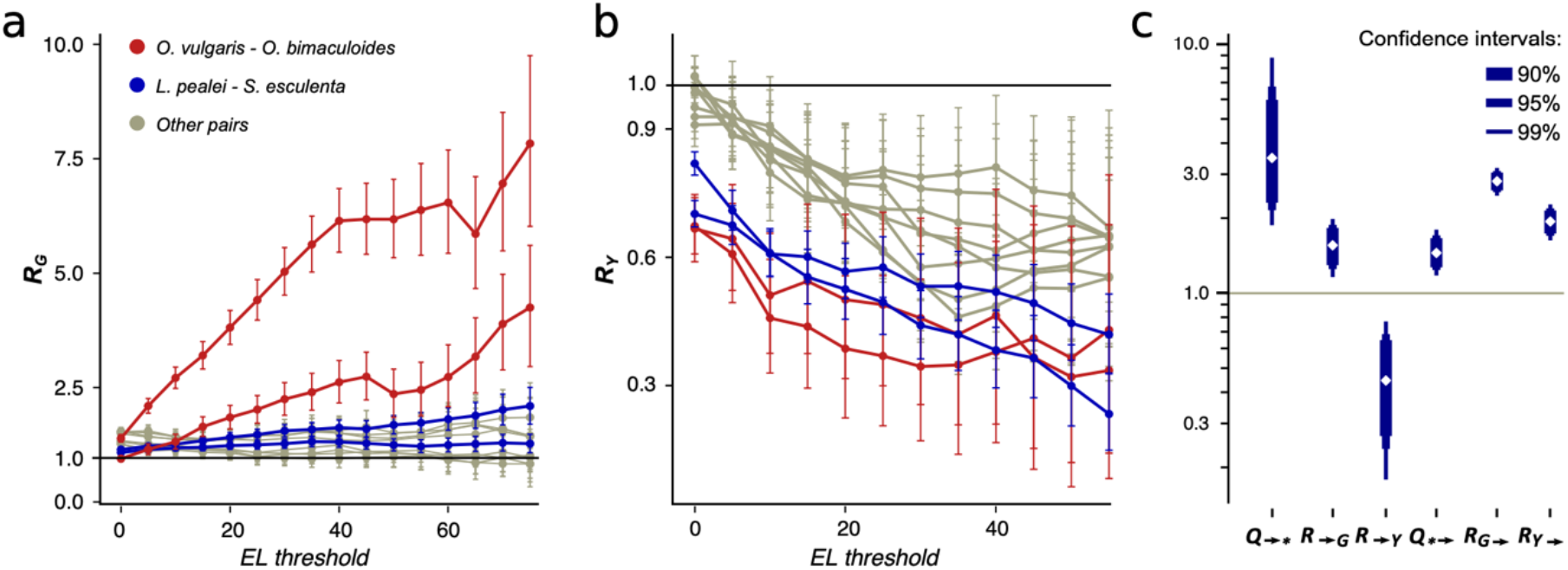
*R* and *Q* values. (a.b) Dependence of *R*_G_ (a) and *R*_Y_ (b) on the editing level. Two curves for each pair are given, since *R_N_* is calculated two times for each pair of species using one of them a a reference each time. The red curves correspond to the pair *O. vulgaris – O. bimaculoides*; the blue curves, to the pair cuttlefish–squid, the grey curves, to distant pairs. **(c)**. Mutational characteristics of editing sites for the squid–cuttlefish summary substitution matrix. Left to right: *Q*_→*_>>1, *R*_→G_>1, *R*_→Y_<1, *Q*_*→_>1, *R*_G→_>>1, *R*_Y→_>1.

We have calculated *R* values separately for non-synonymous editing sites (NES), which comprise between 64.6% and 65.7% of all detected coleoid editing sites, and for synonymous editing sites (SESs) which comprise the remaining 34.3% to 35.4%. NES (Suppl. Fig. S8ab) demonstrate the same pattern as described above for all sites, whereas for SES, we see no dependence of *R*_Y_ on the EL (Suppl. Fig. S8cd). NES demonstrate very low *R*_Y_ at high ELs. These patterns suggest that at highly edited nonsynonymous adenine sites, any nucleotide other than guanines are impeded by strong negative selection; whereas the guanine states at such sites are frequent. Thus, at non-synonymous NCES, the selection patterns differ from those at non-conserved adenines: Y mismatches with NCES experience stronger negative selection than Y mismatches with non-edited adenines, and stronger positive and/or weaker negative selection acting on E-to-G or G-to-E substitutions compared to A-to-G or G-to-A ones, respectively.

### Editing recapitulates substitutions that are positively selected

To reveal the mode of selection at edited sites, we have calculated the *dN/dS* ratios separately for mismatches of edited and unedited adenines with guanines and with pyrimidines (Suppl. Fig. S9). For weakly edited adenines, the *dN/dS* values of mismatches with guanines and with pyrimidines are approximately the same as those for unedited adenines. However, at highly edited sites, the *dN/dS* ratio for substitutions to guanine is two- to threefold higher, compared to unedited adenines, while the respective ratio for pyrimidines is twofold lower. Thus, strongly edited sites evolve under weaker purifying selection against E-to-G and/or G-to-E transitions and stronger purifying selection against E-to-Y and/or Y-to-E substitutions.

To distinguish between positive selection and relaxation of negative selection at these sites, we have calculated *dN/dS* for A-G mismatches where *dN* and *dS* are calculated for edited and unedited adenines, respectively. It is larger than 1 at high ELs for the closely related octopus species pair (Fig. 4), indicating positive selection acting on the E-to-G transition: heavily edited adenines are positively selected for substitutions to guanine.

**Figure 4.**
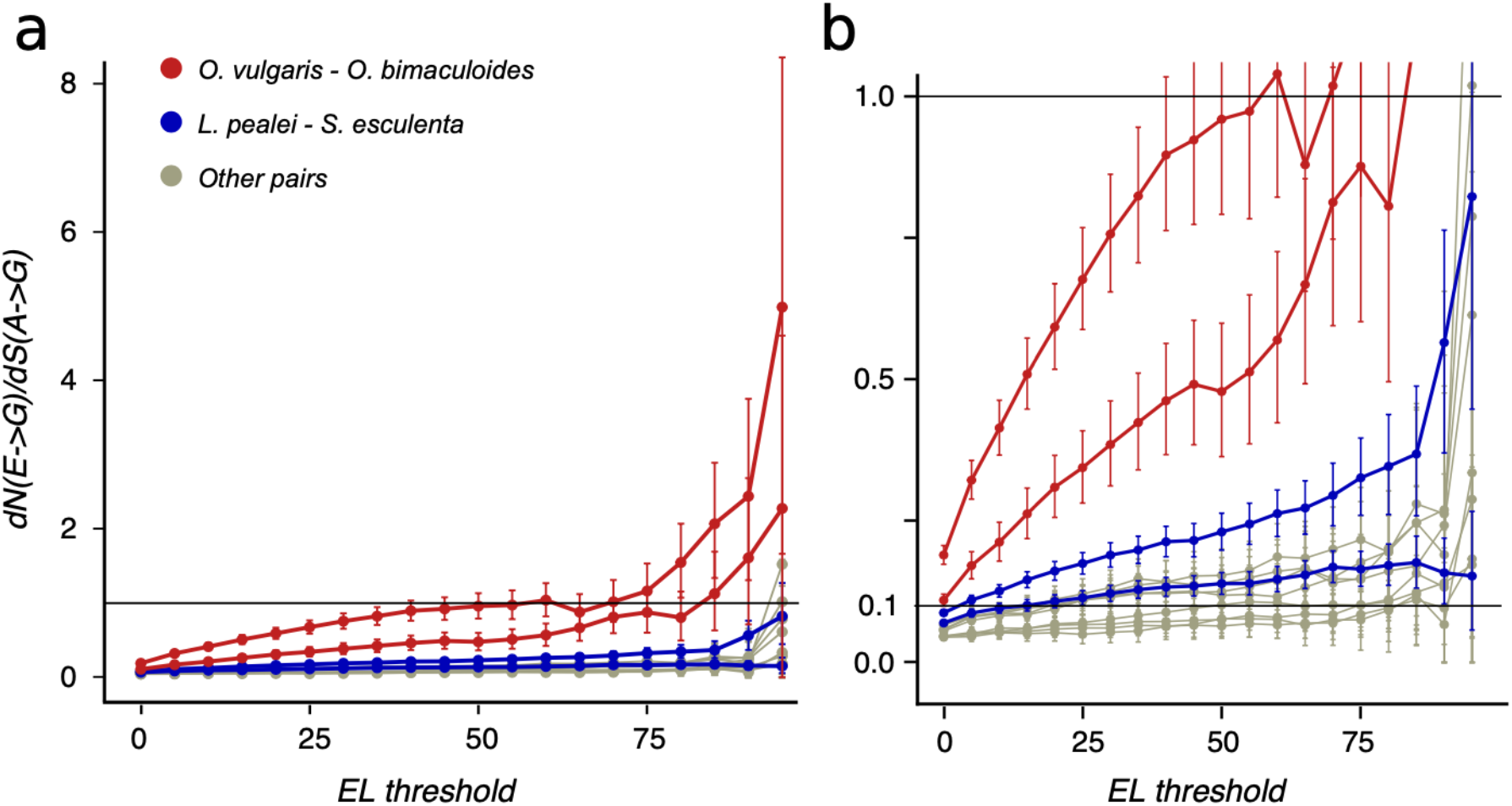
*dN/dS* values of adenine substitutions to guanines for various EL thresholds. Non-synonymous substitutions are calculated for edited adenines, and synonymous, for unedited adenines. Error bars indicate the 95% probability value range. **a**. Plot for the whole range of *dN/dS* values. **b**. Truncated value range. Note the increase of *dN/dS* values at high EL values for all species pairs.

### E-to-G substitutions versus G-to-E substitutions

In theory, two processes could lead to the increase in the observed *R* and *dN/dS* values of edited sites — the increased frequency of either E-to-G or G-to-E substitutions. To distinguish between these possibilities, we use the procedure described in Materials and Methods to сalculate the frequencies of all types of substitutions for each species since its closest ancestor. We also consider the more robust, averaged substitution frequencies for the *Octopus* pair and for the squid–cuttlefish pair. As the frequencies of substitutions to edited and non-edited adenines are calculated separately, we introduce the normalized, directional measure *Q_→*_* reflecting the preference of edited adenines to substitute to guanine:

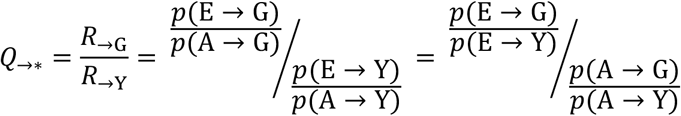

By this definition, the *Q_→*_* measure is an indicator of the joint effect of the prevalence of E-to-G over A-to-G substitutions and of the underrepresentation of E-to-Y relative to A-to-Y substitutions. For the squid–cuttlefish clade, and for SESs and NESs considered separately, *Q_→*_* ranges from 3.49 to 6.4 (Fig. 3c, Suppl. Fig. S10ab), in all cases being significantly higher than 1 expected under a neutral model (*p*<0.005). Hence, as in the case of pairwise comparison of extant species (Fig. 3ab), edited adenines have a substitution pattern strikingly different from that of unedited adenines, and are likely to mutate into guanines.

However, large values of *Q_→*_* may be explained by two effects, high *R*_→G_ of E-to-G substitutions or low *R*_→Y_ of E-to-Y substitutions (Fig. 3c) both yielding *R*_→G_ higher than *R*_→Y_. *R*_→G_ is higher than 1 (*p*<0.005), thus indicating that an edited adenine is more likely to be substituted by guanine than an unedited adenine. Combined with *R*_→Y_ being smaller than 1 (*p*<0.005), this indicates that in fact both effects contribute to the observed *Q*_→*_ values. A similar pattern holds if we consider NES and SES separately: *R*_→G_ is higher than *R*_→Y_, although for NES high *Q*_→*_ can be almost entirely attributed to *R*_→G_, and for SES, to *R*_→Y_ (*p*<0.005) (Suppl. Fig. S10ab).

To analyze the directionality of the mutation process that affects editing states, we consider a similar function measuring the degree of prevalence of G-to-E substitutions:

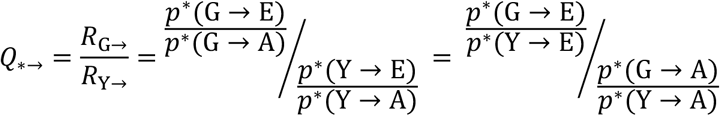

where probabilities *p^*^* are conditional probabilities of a nucleotide mutating to either edited or unedited adenine after taking into account differences in the E and A densities in the transcriptomes:

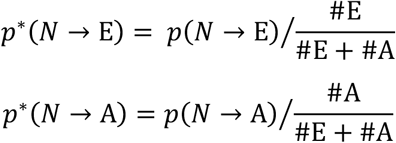

For both the *Octopus* pair and the squid–cuttlefish pair, *Q*_*→_ is larger than 1 (*p*<0.005) (Fig. 3c, Suppl. Fig. S10), thus suggesting that guanines tend to be substituted by edited rather than unedited adenines. However, this effect is on average twofold smaller than that for substitutions of edited adenines to guanines, suggesting that the process of G-to-E transitions is less directional than that for E-to-G transitions. If *R*_G→_ and *R*_Y→_ are considered separately, they both are larger than the expected value 1 (Fig. 3c) (*p*<0.005), which points to a generally faster generation of E sites from both G and Y nucleotides. As the observed effect is small, it could be attributed to weaker negative selection acting upon the G-to-E transition relative to G-to-A, as the edited adenine is a state closer to the guanine-only variant (Jiang and Zhang 2019).

The *Q* value defined as the ratio of undirected *R* values increases with the EL (as follows from Fig. 3ab). On the other hand, formally it is monotonic with respect to the directed *Q_→*_* and *Q*→* values (see Suppl. Mat. 1). Hence, even though we could not detect a significant dependence of *Q_→*_* and *Q_*→_* on EL due to insufficient data, at least one of them should increase with the EL. However, the effects observed for the E-to-G substitution are more pronounced compared with those for the G-to-E substitutions, hinting at A-to-I editing sites mimicking beneficial A-to-G substitutions rather than rescuing deleterious G-to-A substitutions.

## Discussion

### The hypothesis about the adaptivity of non-conserved editing sites is supported by our observations

Editing in coleoids is essential for transcriptome diversification, and results in a more complex phenotype (Liscovitch-Brauer et al. 2017; Eisenberg and Levanon 2018). Indeed, a considerable fraction of coleoid editing sites are conserved between even distantly related species, and a majority of heavily edited sites affect protein sequence (Liscovitch-Brauer et al. 2017). We propose that non-conserved coleoid editing sites could facilitate adaptation by extending selection to regions affecting editing if guanine is the beneficial variant at the editing site. This hypothesis is directly supported by our observations. Indeed, strong dependence of editing on the local context allows for selection of mutations in the vicinity of the editing site, hence extending the variety of beneficial mutations. On the other hand, edited adenines indeed tend to be substituted by guanines, and guanines are selected for if the editing levels of homologous adenines is high. This positive selection pattern is specific to guanine variants, as substitutions of edited adenines to cytosine or thymine are avoided.

An indirect observation also supports our hypothesis. Sizes of the effects such as the E-to-G substitution rate or the rate of positive selection on the guanine variant at editing sites are larger for heavily edited adenines compared to medium and weakly edited ones. This effect could be explained by beneficial A-to-G substitutions provoking selection on adjacent regions, which leads to the increased ELs and hence to the enhanced presence of the guanine-like variant. Indeed, if G is beneficial at a given site, it would manifest as both positive selection towards G at this site, and by mutations at adjacent sites yielding higher A-to-I editing level, and hence these two types of effects would be correlated.

### Positive selection in favor of E-to-G substitutions

Why would substitutions that recapitulate editing be adaptive? Conceivably, it could be that variability at the transcriptome level is advantageous by itself, contributing to the proteome diversity, similar to alternative splicing, alternative transcription and translation starts, *etc* (Raj and van Oudenaarden 2008; Gommans et al. 2009; Pickrell et al. 2010). However, this scenario does not explain positive selection of substitutions to G.

Alternatively, editing might create an unconditionally beneficial variant, so that at an edited site, G is always better than A. Under this scenario, editing could recreate the G allele previously lost due to a deleterious G-to-A mutation, or produce a novel G variant which is favored by selection but has not been present at this site previously (Jiang and Zhang 2019). This scenario is supported by the observed selection favoring guanines at edited sites.

But why would selection in favor of G result in an increased A-to-I editing of a fraction of the transcripts, when a “direct” A-to-G genomic mutation at this site would lead to the same result in 100% of transcripts? One reason could be that mutations creating editing sites and/or increasing editing level are more numerous, and therefore more readily available. For a strongly advantageous mutation (with 4*N*_e_*s* >> 1) that does not preexist in the population, the time till its fixation equals 1/(4*N*_e_*sμ*), where *N*_e_ is the effective population size, *s* is selection in favor of the new mutation, and *μ* is the mutation rate, see eq. 3.22 in Kimura 1983. If two types of mutations can yield the desired phenotype, which of them would be the first to fix in an evolving lineage is determined by the product of the corresponding selection and mutation rates.

Let *μ*_1_ be the rate of the direct mutation, and *s*_1_, selection in its favor. Assume that an increase in the number of favored transcripts can also be achieved by *N* other mutations, each characterized by rate *μ*_2_ and selection *s*_2_. The probability that the editing-enhancing mutation will be the first to occur then equals *Nμ*_2_*s*_2_/(*μ*_1_*s*_1_ *+ Nμ*_2_*s*_2_) (Yampolsky and Stolzfus 2001). If *Nμ*_2_*s*_2_>*μ*_1_*s*_1_, the editing-increasing mutation will typically fix earlier than the direct mutation. As we show, many tens of sites may affect editing, making *N* large, and this scenario likely. For example, if the direct A-to-G substitution confers a 10% increase in fitness, but a 1% increase can be achieved by changes in editing by mutations at each of 20 other sites, then the editing-increasing change will be the first to occur with probability 2/3 if the mutation rates are uniform.

This reasoning only applies if the within-species variability level *N_e_μ* is low (<<1); otherwise each site will carry a preexisting mutation, and the mutation rate will be less relevant (McCandlish and Stoltzfus 2014). Low variability is indeed a characteristic trait of the considered coleoid species, with synonymous-site pairwise divergence of 2.5×10^−3^ for *O. bimaculoides*, 2.2×10^−3^ for *O. vulgaris*, 1.8×10^−3^ for *S. eculenta*, and 4.5×10^−3^ for *L. pealei* (see Materials and Methods). These values imply *N_e_μ*<<1, suggesting that evolution can be indeed mutation-limited in this group of species. Low values of *N_e_μ* characteristic of higher animals have been proposed to underlie many aspects of genomic complexity (Lynch 2007); they may also cause the high prevalence of RNA editing in coleoids.

When this study had been completed, Popitsch et al. published a population-genetic study of *Drosophila* and human A-to-I RNA editing sites, in which they showed a similar pattern of selection at editing sites, with the derived G state selected upon, whereas C and T variants being suppressed, indicating enhanced negative selection (Popitsch et al. 2020). That study indirectly supports our claim about coleoid A-to-I editing sites mimicking beneficial A-to-G substitutions. Furthermore, as coleoids possess many more conserved re-coding A-to-I editing sites than any other studied metazoan group, one might expect the bulk of coleoid editing, especially at heavily edited sites, to be important *per se*, *e.g.* for transcriptome diversification, which would result in suppression of any non-adenine variants in editing sites. On the contrary, we have observed positive selection in A-G mismatches, when adenines are heavily edited, with selection acting specifically on A-to-G transitions. Also, like Popitsch et al., we have observed enhanced negative selection against A-to-C and A-to-T substitutions and mismatches at coleoid editing sites. The consistency of results obtained for coleoids, *Drosophila*, and human points towards a general role of A-to-I editing sites as imitations and precursors of A-to-G transitions in the evolution of metazoans with low-polymorphic populations.

### Conservation and function of editing

Earlier, it has been proposed that most editing sites result from tolerable promiscuous ADAR action (Xu and Zhang 2014). However, the A-to-I editing sites in coleoids are under positive selection if ELs are high (Fig. 4). Hence large ELs cannot result simply from the tolerance towards substitutions to guanines at these sites.

Сoleoid editing sites are often considered to be important for complex regulation (Albertin et al. 2015; Alon et al. 2015; Liscovitch-Brauer et al. 2017; Eisenberg and Levanon 2018; Jiang and Zhang 2019). However, this applies only to conserved, and hence functional, editing sites. We propose that coleoid editing sites form two populations with different properties. Firstly, there are functional editing sites, which are important *per se* due to their ability to diversify protein products in various tissues and environmental conditions (Savva et al. 2012; Alon et al. 2015; Harjanto et al. 2016; Buchumenski et al. 2017; Duan et al. 2017; Liscovitch-Brauer et al. 2017; Tan et al. 2017). As such sites should be retained over long periods of time, we may consider conservation as a good proxy for functionality. Conserved sites are surrounded by conserved regions (Liscovitch-Brauer et al. 2017), their ELs show dependence on the number of substitutions in adjacent regions (Suppl. Fig. S4), and they comprise up to about a half of A-to-I editing sites in a coleoid transcriptome (Liscovitch-Brauer et al. 2017).

Secondly, there are non-functional sites; the proxy here are non-conserved sites, with a caveat that some recently emerged sites could be functional. Nonetheless, as the proportion of young functional sites should be minimal (Gommans et al. 2009), the general properties of non-conserved sites should reasonably well represent those of non-functional ones. Non-conserved sites are not flanked by conserved regions, their ELs show no correlation with the number of substitutions in adjacent regions, and their sequence contexts differ from those of the conserved ones (Suppl. Fig. S2). Our hypothesis that (non-conserved) editing sites have an intrinsic evolutionary value does not contradict the fact that some (possibly large) subset of editing sites are functional as editing sites *per se* from the physiological point of view.

Theoretically, our results could have been influenced by underprediction of editing sites. As the mean EL is about 5%, a site might be easily missed especially in transcripts with low expression levels (Bahn et al. 2011; Alon et al. 2012; Liscovitch-Brauer et al. 2017). However, the majority of our observations are obtained for highly edited adenines, which are predicted with greater accuracy (Bahn et al. 2011), and hence should not be influenced by missing weakly edited sites.

### Theoretical frameworks and alternative explanations

Our results could be interpreted within several paradigms. Firstly, as noted above, the observations could mean that editing rescues deleterious G-to-A substitutions (Jiang and Zhang 2019). However, as also mentioned above, the estimates of *Q* values, which represent the mutation process directionality, indicate that E-to-G substitutions differ in terms of the transition/transversion rate from A-to-G ones to a much greater extent, than G-to-E substitutions differ from G-to-A (*Q*_→*_ >> *Q*_*→_,); in addition, *Q*_*→_<1 at non-synonymous sites (Fig. 3c), again supporting the idea that the E-to-G transitions contribute to the observed effects to a larger degree. Ultimately, this issue would be resolved when more data are available, allowing for the reconstruction of ancestral states of NCES.

Our results could be formulated in terms of Waddington’s Genetic Assimilation (Waddington 1953a, 1953b; Lynch and Walsh 1998; Crispo 2007; Ghalambor et al. 2007; Ghalambor et al. 2015; Levis and Pfennig 2016; Ho and Zhang 2018; Levis and Pfennig 2019). Editing could buffer coleoids against environmental changes — under novel conditions the phenotype changes (adenine is edited and read as guanine), and subsequently this change is reinforced on the genome level by the selection process, which we observe as positive selection pressure on E-to-G transitions. However, at present we have no data on environmental variance in the coleoid A-to-I editing.

The preadaptation paradigm refers to a pre-existing structure that has changed its function or acquired a new one in the course of evolution (Darwin 1872; Gould and Vrba 1982; McLennan 2008; Ardila 2016; Casinos 2017; Cadotte et al. 2018). Here, as non-functional editing should be mostly effectively neutral (Gommans et al. 2009), it might generate a pool of variants, some of which may become advantageous in the future, when the genetic background or environmental conditions change. However, to claim preadaptation one should determine the function of each editing site, which is not feasible.

Hence, the most reasonable framework for our findings seems to be in terms of non-functional editing sites enhancing the expressed genetic variability, thus contributing to the acceleration of the evolutionary process at sites with beneficial A-to-G substitution. The Continuous Probing Hypothesis (Gommans et al. 2009) states that editing sites, due to the lack of a strict context, constantly emerge at random points of the transcribed genomic regions. Hence, an adenine with a beneficial substitution to guanine could become edited if the editing context emerges around it purely by chance. The context can be further selected upon, resulting in the mimicking of the beneficial guanine variant. (An extended version of this discussion is provided as Suppl. Mat. 3)

A similar rhetoric can be applied to other cellular information transmission processes such as transcription and splicing. These processes depend on regulatory sites and contexts that change the quantity, dynamics (developmental stage, tissue-specificity, response to external conditions), and sequence of encoded proteins and hence are subject to selection (Raj and van Oudenaarden 2008; Pickrell et al. 2010). Hence a natural extension of this study would be to systematically assess the evolutionary advantage of noise in information transmission processes in low-polymorphic populations.

## Materials and Methods

### Data

Transcriptomes for all six considered species, *O. vulgaris*, *O. bimaculoides*, *S. esculenta*, *L. pealei*, *N. pompilius*, and *A. californica*, parameters of editing sites, and tables of conserved editing sites were taken from the online supplementary data of Liscovitch-Brauer et al. 2017. Genomic read data were downloaded from the SRA database. *S. esculenta* and *O. vulgaris* genomic read data were taken from bioproject PRJNA299756, *L. pealei*, from PRJNA255916, and *O. bimaculoides*, from PRJNA270931.

### Annotation of structured and unstructured regions

To estimate the structural potential of each position we used Z-score values obtained by the RNASurface program (Soldatov et al. 2013). Here, Z-score of a sequence is defined as *Z* = (E - μ)/σ where *E* is the minimal free energy of a biological sequence, μ and σ are the mean and standard deviation of the energy distribution of shuffled sequences with preserved length and average dinucleotide composition. The program was run with parameters maximal sliding window length 350 and minimal sliding window length 20. From the RNASurface output, structural potential of overlapping segments was inferred. Each position of each transcript was assigned the best (minimal) *Z-score* of all structured segments containing it, if it was less than –2, otherwise it was assigned null value. As a result, each transcript was divided into structured and unstructured regions with a *Z-*score value assigned to all positions in the structured regions. The difference between the structural potential upon the A-to-G change (Fig. 2d) was considered if its absolute value exceeded 2.

### Analysis of polymorphisms

Genomic reads were mapped onto transcriptomes with bowtie2 (Langmead and Salzberg 2012) using the --sensitive-local run mode. After the sorting of the resulting read alignment files with the samtools package (Li et al. 2009), diploid genotypes were called with bcftools (Narasimhan et al. 2016). Next, we discarded all non-SNP variants and variants with the quality score below 20. We computed synonymous nucleotide diversity πs with the pairwise haplotype comparison implemented in the PAML package (Yang 2007).

### Alignments

To construct multiple transcriptome alignments, we selected a transcriptome of one species and performed BLASTn (Altschul et al. 1990) with the E-value threshold of 10^−15^ against the transcriptomes of the remaining species. Resulting alignment was obtained by merging of the pairwise BLASTn alignments. The results showed only a negligible dependence on the choice of the seed species.

### Context analysis

Site LOGOs were built with the WebLOGO server (Crooks 2004). *R* values for mismatches in contexts of non-conserved editing sites were defined as:

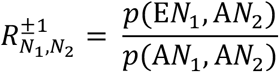

where *N_1_* and *N_2_* represent nucleotides in positions +1 and –1 relative to the considered adenine, *p*(E*N_1_*, A*N_2_*) is the probability of a mismatch in the position +1 or –1 relative to the considered adenine that is edited in one of the two considered species and not edited an another, and *p*(A*N_1_*, A*N_2_*) is the respective probability in the case when both homologous adenines are non-edited. The statistical significance of the *R* values was assessed by the chi-squared contingency test with the Bonferroni correction.

### Substitution matrix

For a considered species, we considered its closest relative and an outgroup that could be either of the two remaining coleoids (Fig. 1a). Given the low number of available species, we used maximum parsimony (MP) to reconstruct ancestral states. Thus, for a position in the alignment, the ancestral state of nucleotide *N* was inferred if the closest relative and an outgroup had the same nucleotide *N*^anc^; an ancestral adenine was considered to be edited if the homologous adenines in the closest relative and an outgroup were edited. The substitution matrix was thus comprised of counts inferred by MP, #(*N*^anc^ → *N*).

### *R* and *Q* calculation for non-synonymous (NES) and synonymous (SES) editing sites

When *R* measures were computed separately for SES and NES, we applied a modification of the expression for the *R* value. For substitutions at SES:

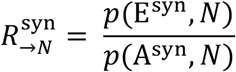

where E^syn^ are synonymous editing sites, i.e. edited adenines that, when substituted to guanine, do not change the amino acid, and, similarly, A^syn^ are synonymous unedited adenines. An analogous formula was applied for non-synonymous editing sites.

When we calculated *R*_*N*→_ separately for NES and SES, we applied another modification of the expression for the *R* value. For mutations to SES we have:

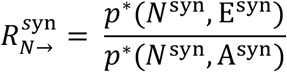

where E^syn^ and A^syn^ are defined just as above, and *N*^syn^ represents nucleotides, that, when substituted with adenine and with guanine would yield the same amino acid. *p*∗ are, just like in formula (3), conditional probabilities:

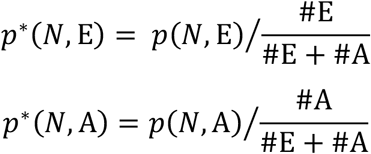

An analogous formula is applied to NES, with *N*^non^ representing nucleotides, that, when substituted with adenine and with guanine would yield different amino acids.

### Calculation of *dN/dS*

We estimated the strength of positive selection acting on substitutions to guanines and to pyrimidines separately by applying the *dN/dS* measure to edited adenines with a subsequent normalization by *dN/dS* of unedited adenines. Thus, for substitutions to G we applied the formula (Suppl. Fig. S11):

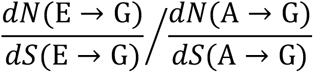

where *dN* were calculated for all codons and *dS*, for four- and six-fold degenerate codons. An analogous formula was used to estimate selection acting on E-to-Y substitutions. Next, we applied this measure separately for 10% EL bins, counted Pearson’s correlation coefficient and applied the *F* statistic to estimate the significance of the obtained correlation.

Positive selection on editing sites was estimated with the *dN/dS* ratio where non-synonymous substitutions were considered for edited adenines, and synonymous, for unedited adenines:

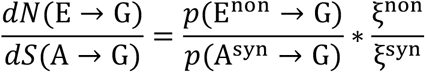

where ξ^non^ and ξ^syn^ are normalizing coefficients accounting for differences in codon probabilities and different probabilities of, respectively, synonymous and non-synonymous substitutions under the neutral evolution assumption. These coefficients are defined as:

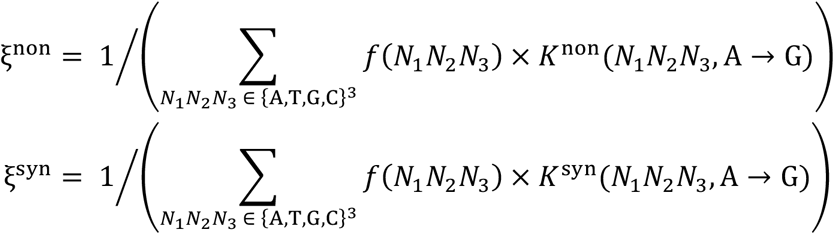

where *f(N_1_N_2_N_3_)* is the codon frequency while *K*^non^ and *K*^syn^ are, respectively, the numbers of possible non-synonymous and synonymous A-to-G substitutions in a given codon.

### Statistics

For mutation frequency and *dN/dS* analysis, statistics were obtained from 10^5^ random sets of mutation numbers sampled from the binomial distributions with the parameters equal to the observed substitution frequencies. For the analysis of parallel evolution, two-tailed confidence intervals were inferred from the binomial distribution. The binomial test was applied to compare fractions of conserved and not conserved editing sites in structured segments. For the analysis of changes in the secondary structure stability following A-to-G *in silico* substitutions, sizes of tails in the distribution of Z-score differences were compared using the binomial test, and to compare the results for different types of sites, random 100-sequence samples of each type were compared with the Wilcoxon signed-rank test.

### Code availability

*Ad hoc* scripts were written in Python. Graphs were built using R. All scripts and data analysis protocols are available online at https://github.com/mikemoldovan/coleoidRNAediting.

## Supporting information

File containing all supplementary data

## Funding

This study was supported by the Russian Foundation of Basic Research under grant 18-29-13011

The authors are grateful to Stepan Denisov, Sofya Garushyants, Fyodor Kondrashov, and Eugene Koonin for discussions, criticisms, and suggestions.

## Notes

### Competing Interest Statement

The authors have declared no competing interest.

## References

Albertin CB, Simakov O, Mitros T, Wang ZY, Pungor JR, Edsinger-Gonzales E, Brenner S, Ragsdale CW, Rokhsar DS. 2015. The octopus genome and the evolution of cephalopod neural and morphological novelties. Nature 524, 220–224. (doi:10.1038/nature14668)

Alon S, Garrett SC, Levanon EY, Olson S, Graveley BR, Rosenthal JJC, Eisenberg E. 2015. The majority of transcripts in the squid nervous system are extensively recoded by A-to-I RNA editing. eLife 4. (doi:10.7554/elife.05198)

Alon S, Mor E, Vigneault F, Church GM, Locatelli F, Galeano F, Gallo A, Shomron N, Eisenberg E. 2012. Systematic identification of edited microRNAs in the human brain. Genome Research 22, 1533–1540. (doi:10.1101/gr.131573.111)

Altschul SF, Gish W, Miller W, Myers EW, Lipman DJ. 1990. Basic local alignment search tool. Journal of Molecular Biology 215, 403–410. (doi:10.1016/s0022-2836(05)80360-2)

Ardila A. 2016. The Evolutionary Concept of “Preadaptation” Applied to Cognitive Neurosciences. Frontiers in Neuroscience 10. (doi:10.3389/fnins.2016.00103)

Avery PJ, Hill WG. 1977. Variability in genetic parameters among small populations. Genetical Research 29, 193–213. (doi:10.1017/s0016672300017286)

Bahn JH, Lee J-H, Li G, Greer C, Peng G, Xiao X. 2011. Accurate identification of A-to-I RNA editing in human by transcriptome sequencing. Genome Research 22, 142–150. (doi:10.1101/gr.124107.111)

Barton N, Partridge L. 2000. Limits to natural selection. BioEssays 22, 1075–1084. (doi:10.1002/1521-1878(200012)22:12<1075::aid-bies5>3.0.co;2-m)

Bass BL, Weintraub H. 1988. An unwinding activity that covalently modifies its double-stranded RNA substrate. Cell 55, 1089–1098. (doi:10.1016/0092-8674(88)90253-x)

Buchumenski I, Bartok O, Ashwal-Fluss R, Pandey V, Porath HT, Levanon EY, Kadener S. 2017. Dynamic hyper-editing underlies temperature adaptation in Drosophila. PLOS Genetics 13, e1006931. (doi:10.1371/journal.pgen.1006931)

Cadotte MW, Campbell SE, Li S, Sodhi DS, Mandrak NE. 2018. Preadaptation and Naturalization of Nonnative Species: Darwin’s Two Fundamental Insights into Species Invasion. Annual Review of Plant Biology 69, 661–684. (doi:10.1146/annurev-arplant-042817-040339)

Casinos A. 2017. From Cuénot’s préadaptation to Gould and Vrba’s exaptation: a review. Biological Journal of the Linnean Society 121, 239–247. (doi:10.1093/biolinnean/blw038)

Crispo E. 2007. The Baldwin Effect and Genetic Assimilation: Revisiting two Mechanisms of Evolutionary Change Mediated by Phenotypic Plasticity. Evolution 61, 2469–2479. (doi:10.1111/j.1558-5646.2007.00203.x)

Crooks GE. 2004. WebLogo: A Sequence Logo Generator. Genome Research 14, 1188–1190. (doi:10.1101/gr.849004)

Darwin, CR. 1872. The origin of species by means of natural selection, or the preservation of favoured races in the struggle for life, 6th edition; with additions and corrections. London: John Murray.

Duan Y, Dou S, Luo S, Zhang H, Lu J. 2017. Adaptation of A-to-I RNA editing in Drosophila. PLOS Genetics 13, e1006648. (doi:10.1371/journal.pgen.1006648)

Durrett R, Schmidt D. 2008. Waiting for Two Mutations: With Applications to Regulatory Sequence Evolution and the Limits of Darwinian Evolution. Genetics 180, 1501–1509. (doi:10.1534/genetics.107.082610)

Eisenberg E, Levanon EY. 2018. A-to-I RNA editing — immune protector and transcriptome diversifier. Nature Reviews Genetics 19, 473–490. (doi:10.1038/s41576-018-0006-1)

Ensterö M, Daniel C, Wahlstedt H, Major F, Öhman M. 2009. Recognition and coupling of A-to-I edited sites are determined by the tertiary structure of the RNA. Nucleic Acids Research 37, 6916–6926. (doi:10.1093/nar/gkp731)

Farajollahi S, Maas S. 2010. Molecular diversity through RNA editing: a balancing act. Trends in Genetics 26, 221–230. (doi:10.1016/j.tig.2010.02.001)

Ghalambor CK, Hoke KL, Ruell EW, Fischer EK, Reznick DN, Hughes KA. 2015. Non-adaptive plasticity potentiates rapid adaptive evolution of gene expression in nature. Nature 525, 372–375. (doi:10.1038/nature15256)

Ghalambor CK, McKay JK, Carroll SP, Reznick DN. 2007. Adaptive versus non-adaptive phenotypic plasticity and the potential for contemporary adaptation in new environments. Functional Ecology 21, 394–407. (doi:10.1111/j.1365-2435.2007.01283.x)

Gommans WM, Mullen SP, Maas S. 2009. RNA editing: a driving force for adaptive evolution? BioEssays 31, 1137–1145. (doi:10.1002/bies.200900045)

Gould SJ, Vrba ES. 1982. Exaptation—a Missing Term in the Science of Form. Paleobiology 8, 4–15. (doi:10.1017/s0094837300004310)

Harjanto D, Papamarkou T, Oates CJ, Rayon-Estrada V, Papavasiliou FN, Papavasiliou A. 2016. RNA editing generates cellular subsets with diverse sequence within populations. Nature Communications 7. (doi:10.1038/ncomms12145)

Hedges SB, Dudley J, Kumar S. 2006. TimeTree: a public knowledge-base of divergence times among organisms. Bioinformatics 22, 2971–2972. (doi:10.1093/bioinformatics/btl505)

Ho W-C, Zhang J. 2018. Evolutionary adaptations to new environments generally reverse plastic phenotypic changes. Nature Communications 9. (doi:10.1038/s41467-017-02724-5)

Jiang D, Zhang J. 2019. The preponderance of nonsynonymous A-to-I RNA editing in coleoids is nonadaptive. Nature Communications 10. (doi:10.1038/s41467-019-13275-2)

Jin Y, Zhang W, Li Q. 2009. Origins and evolution of ADAR-mediated RNA editing. IUBMB Life 61, 572–578. (doi:10.1002/iub.207)

Kim DDY. 2004. Widespread RNA Editing of Embedded Alu Elements in the Human Transcriptome. Genome Research 14, 1719–1725. (doi:10.1101/gr.2855504)

Kimura M. 1983. The Neutral Theory of Molecular Evolution. Cambridge University Press.

Klironomos FD, Berg J, Collins S. 2013. How epigenetic mutations can affect genetic evolution: Model and mechanism. BioEssays 35, 571–578. (doi:10.1002/bies.201200169)

Kronholm I, Collins S. 2015. Epigenetic mutations can both help and hinder adaptive evolution. Molecular Ecology 25, 1856–1868. (doi:10.1111/mec.13296)

Kurmangaliyev YZ, Ali S, Nuzhdin SV. 2015. Genetic Determinants of RNA Editing Levels of ADAR Targets in Drosophila melanogaster. G3: Genes|Genomes|Genetics 6, 391–396. (doi:10.1534/g3.115.024471)

Lanfear R, Kokko H, Eyre-Walker A. 2014. Population size and the rate of evolution. Trends in Ecology & Evolution 29, 33–41. (doi:10.1016/j.tree.2013.09.009)

Langmead B, Salzberg SL. 2012. Fast gapped-read alignment with Bowtie 2. Nature Methods 9, 357–359. (doi:10.1038/nmeth.1923)

Levis NA, Pfennig DW. 2016. Evaluating “Plasticity-First” Evolution in Nature: Key Criteria and Empirical Approaches. Trends in Ecology & Evolution 31, 563–574. (doi:10.1016/j.tree.2016.03.012)

Levis NA, Pfennig DW. 2019. Phenotypic plasticity, canalization, and the origins of novelty: Evidence and mechanisms from amphibians. Seminars in Cell & Developmental Biology 88, 80–90. (doi:10.1016/j.semcdb.2018.01.012)

Lewontin RC. 1964. The Interaction of Selection and Linkage. General Considerations; Heterotic Models. Genetics. 49(1):49–67.

Li H et al. 2009. The Sequence Alignment/Map format and SAMtools. Bioinformatics 25, 2078–2079. (doi:10.1093/bioinformatics/btp352)

Liscovitch-Brauer N et al. 2017. Trade-off between Transcriptome Plasticity and Genome Evolution in Cephalopods. Cell 169, 191–202.e11. (doi:10.1016/j.cell.2017.03.025)

Lush, JL. 1937. Animal Breeding Plans. Ames, Iowa: Iowa State Press.

Lynch M. 2007. The Origins of Genome Architecture. Sinauer Associates.

Lynch, M. & Walsh, B. 1998. Genetics and Analysis of Quantitative Traits, 1st edn. Sinauer, Sunderland, MA.

McCandlish DM, Stoltzfus A. 2014. Modeling Evolution Using the Probability of Fixation: History and Implications. The Quarterly Review of Biology 89, 225–252. (doi:10.1086/677571)

McLennan DA. 2008. The Concept of Co-option: Why Evolution Often Looks Miraculous. Evolution: Education and Outreach 1, 247–258. (doi:10.1007/s12052-008-0053-8)

Morse DP, Aruscavage PJ, Bass BL. 2002. RNA hairpins in noncoding regions of human brain and Caenorhabditis elegans mRNA are edited by adenosine deaminases that act on RNA. Proceedings of the National Academy of Sciences 99, 7906–7911. (doi:10.1073/pnas.112704299)

Nam K, Munch K, Mailund T, Nater A, Greminger MP, Krützen M, Marquès-Bonet T, Schierup MH. 2017. Evidence that the rate of strong selective sweeps increases with population size in the great apes. Proceedings of the National Academy of Sciences 114, 1613–1618. (doi:10.1073/pnas.1605660114)

Narasimhan V, Danecek P, Scally A, Xue Y, Tyler-Smith C, Durbin R. 2016. BCFtools/RoH: a hidden Markov model approach for detecting autozygosity from next-generation sequencing data. Bioinformatics 32, 1749–1751. (doi:10.1093/bioinformatics/btw044)

Pickrell JK, Pai AA, Gilad Y, Pritchard JK. 2010. Noisy Splicing Drives mRNA Isoform Diversity in Human Cells. PLoS Genetics 6, e1001236. (doi:10.1371/journal.pgen.1001236)

Pinto Y, Cohen HY, Levanon EY. 2014. Mammalian conserved ADAR targets comprise only a small fragment of the human editosome. Genome Biology 15, R5. (doi:10.1186/gb-2014-15-1-r5)

Popitsch N, Huber CD, Buchumenski I, Eisenberg E, Jantsch M, von Haeseler A, Gallach M. 2020. A-to-I RNA Editing Uncovers Hidden Signals of Adaptive Genome Evolution in Animals. Genome Biology and Evolution 12:345–357. (doi:10.1093/gbe/evaa046)

Raj A, van Oudenaarden A. 2008. Nature, Nurture, or Chance: Stochastic Gene Expression and Its Consequences. Cell 135, 216–226. (doi:10.1016/j.cell.2008.09.050)

Ramaswami G, Lin W, Piskol R, Tan MH, Davis C, Li JB. 2012. Accurate identification of human Alu and non-Alu RNA editing sites. Nature Methods 9, 579–581. (doi:10.1038/nmeth.1982)

Reenan RA. 2005. Molecular determinants and guided evolution of species-specific RNA editing. Nature 434, 409–413. (doi:10.1038/nature03364)

Rieder LE, Staber CJ, Hoopengardner B, Reenan RA. 2013. Tertiary structural elements determine the extent and specificity of messenger RNA editing. Nature Communications 4. (doi:10.1038/ncomms3232)

Rousselle M, Simion P, Tilak M-K, Figuet E, Nabholz B, Galtier N. 2020. Is adaptation limited by mutation? A timescale-dependent effect of genetic diversity on the adaptive substitution rate in animals. PLOS Genetics 16, e1008668. (doi:10.1371/journal.pgen.1008668)

Savva YA, Rieder LE, Reenan RA. 2012. The ADAR protein family. Genome Biology 13, 252. (doi:10.1186/gb-2012-13-12-252)

Smith JM. 1976. What Determines the Rate of Evolution? The American Naturalist. 110(973):331–8.

Soldatov RA, Vinogradova SV, Mironov AA. 2013. RNASurface: fast and accurate detection of locally optimal potentially structured RNA segments. Bioinformatics 30, 457–463. (doi:10.1093/bioinformatics/btt701)

Tan MH et al. 2017. Dynamic landscape and regulation of RNA editing in mammals. Nature 550, 249–254. (doi:10.1038/nature24041)

Waddington CH. 1953a. Genetic Assimilation of an Acquired Character. Evolution 7, 118–126. (doi:10.1111/j.1558-5646.1953.tb00070.x)

Waddington CH. 1953b. The Baldwin Effect, Genetic Assimilation AND Homeostasis. Evolution 7, 386–387. (doi:10.1111/j.1558-5646.1953.tb00099.x)

Wang AH-J, Hakoshima T, van der Marel G, van Boom JH, Rich A. 1984. AT base pairs are less stable than GC base pairs in Z-DNA: The crystal structure of d(m5CGTAm5CG). Cell 37, 321–331. (doi:10.1016/0092-8674(84)90328-3)

Xu G, Zhang J. 2014. Human coding RNA editing is generally nonadaptive. Proceedings of the National Academy of Sciences 111, 3769–3774. (doi:10.1073/pnas.1321745111)

Yablonovitch AL, Deng P, Jacobson D, Li JB. 2017. The evolution and adaptation of A-to-I RNA editing. PLOS Genetics 13, e1007064. (doi:10.1371/journal.pgen.1007064)

Yampolsky LY, Stoltzfus A. 2001. Bias in the introduction of variation as an orienting factor in evolution. Evolution and Development 3, 73–83. (doi:10.1046/j.1525-142x.2001.003002073.x)

Yang Y, Lv J, Gui B, Yin H, Wu X, Zhang Y, Jin Y. 2008. A-to-I RNA editing alters less-conserved residues of highly conserved coding regions: Implications for dual functions in evolution. RNA 14, 1516–1525. (doi:10.1261/rna.1063708)

Yang Z. 2007. PAML 4: Phylogenetic Analysis by Maximum Likelihood. Molecular Biology and Evolution 24, 1586–1591. (doi:10.1093/molbev/msm088)

Yu Y, Zhou H, Kong Y, Pan B, Chen L, Wang H, Hao P, Li X. 2016. The Landscape of A-to-I RNA Editome Is Shaped by Both Positive and Purifying Selection. PLOS Genetics 12, e1006191. (doi:10.1371/journal.pgen.1006191)

