## Supplementary material for "Adaptive evolution at mRNA editing sites in soft-bodied cephalopods": File containing all supplementary data

### Supplementary materials for the paper "Adaptive evolution at mRNA editing sites in soft-bodied cephalopods"

#### 1. Extrapolation

Unpolarized *R* values strongly depend on the EL threshold. Due to insufficient data, analogous, statistically significant observations for polarized *Q* values at EL thresholds higher than 0 could not be obtained; however, we can indirectly demonstrate that *Q* must vary at different EL levels. Consider two closely related organisms, "1" and "2", and their common ancestor, "anc". By definition given in the main text:

$$Q^{1\to 2} = \frac{R^{1\to 2}_{\to \mathrm{G}}}{R^{1\to 2}_{\to \mathrm{Y}}} = \left.\frac{p(\mathrm{E}^1 \to \mathrm{G}^2)}{p(\mathrm{A}^1 \to \mathrm{G}^2)}\right/ \frac{p(\mathrm{E}^1 \to \mathrm{Y}^2)}{p(\mathrm{A}^1 \to \mathrm{Y}^2)} =$$

$$= \frac{\left(p(\mathrm{E}^{\mathrm{anc}} \to \mathrm{G}^2) + p(\mathrm{G}^{\mathrm{anc}} \to \mathrm{E}^1)\right) \times \left(p(\mathrm{A}^{\mathrm{anc}} \to \mathrm{Y}^2) + p(\mathrm{Y}^{\mathrm{anc}} \to \mathrm{A}^1)\right)}{\left(p(\mathrm{A}^{\mathrm{anc}} \to \mathrm{G}^2) + p(\mathrm{G}^{\mathrm{anc}} \to \mathrm{A}^1)\right) \times \left(p(\mathrm{E}^{\mathrm{anc}} \to \mathrm{Y}^2) + p(\mathrm{Y}^{\mathrm{anc}} \to \mathrm{E}^1)\right)}$$

Thus, by definition, $Q^{1\to 2}$ is monotonic on both $Q^2_{\to *}$ and $Q^1_{*\to}$. Similarly, $Q^{2\to 1}$ is monotonic on both $Q^1_{\to *}$ and $Q^2_{*\to}$

As no other terms are present in the ratio of *R* values, we conclude that at least one of the directed *Q* values should increase at the increase of the *R* ratio, the latter being observed in the case of its dependence on the EL threshold. Hence at higher EL values, at least one directed *Q* value should increase.

#### 2. Discussion in terms of Genetic Assimilation and Preadaptation

Conrad Hal Waddington has proposed that a trait exhibiting extreme values in a novel environment due to the phenotypic variation present in a population will be canalized through subsequent evolution (Waddington 1953a, 1953b). Evidently, the trait has to be phenotypically plastic, i.e. to exhibit environmental variation and/or genotype×environment interaction covariance (Lynch and Walsh 1998; Ghalambor et al. 2015; Levis and Pfennig 2019). Over the years, the subject of phenotypic plasticity generally facilitating adaptation through genetic assimilation remained debatable (Ghalambor et al. 2015), and, despite multiple examples of genetic assimilation (Levis and Pfennig 2019), large-scale studies of differential expression of genes under environmental changes before introduction to a novel environment and in the novel environment before and after adaptation have shown that genetic changes tend to reverse rather than enhance the plastic ones (Ghalambor et al. 2015). Considering that editing may be influenced by environmental changes, one may imagine the environmental variance of editing, and, moreover, directed changes in the editing status in novel environments (Duan et al. 2017). Thus, positive selection acting on E-to-G transitions may be interpreted as genetic changes reinforcing phenotypic changes in the course of adaptation. However, the reinforcement of plastic changes by subsequent adaptation is notoriously difficult to prove, and the research design has to meet a number of specific criteria, such as the ability to show the very presence of environmental variance, which we do not have sufficient data to test (Duan et al. 2017).

This explanation is also contradicted by A-to-I editing being performed by a single small family of ADAR enzymes (Eisenberg and Levanon, 2018), that for simple combinatorial reasons cannot provide a complex and specific response to a novel environment as differential expression. Nonetheless, editing in coleoids could be regulated by other proteins or processes such as the dependence of local RNA structures on temperature, and hence coleoid mRNA editing could be a good object for future studies of genetic assimilation.

Yet another possible explanation is in terms of preadaptation. The term «preadaptation» refers to a pre-existing structure that has changed its function in the course of evolution, a concept introduced by Charles Darwin (Gould and Vrba 1982; McLennan 2008), and currently it is applied to both morphological and molecular traits. This definition presumes that we can identify three stages of the structure's evolution: (1) structure with the ancestral function, (2) structure that has acquired a novel, derived function, but retained the ancestral one, and (3) structure with only the derived function (Gould and Vrba 1982; McLennan 2008). Stage (1) is optional, as a structure that would be beneficial in the future could emerge by neutral evolution and have no specific ancestral function (McLennan 2008). In the case of non-conserved editing we seem to observe a rather similar pattern — an ancestral adenine that, through a transitory stage of an edited nucleotide, where two mRNA isoforms are present, is substituted with guanine. Under this hypothesis, we would expect edited adenines to be substituted more frequently and directionally to guanines, positive selection to act on such substitutions, and these effects to be more pronounced for highly edited sites and for more closely related species, as preadaptation has been shown to be better seen when closely related species are considered. These criteria are basically the same as those listed in the Introduction section, and hence the preadaptation scenario could be applied here. One problem is, that *de facto* we have not observed the complete chain of events, our findings being restricted only to finding that E-to-G transitions are selected for and to frequent emergence of editing sites from unedited adenines. Another problem with this explanation is that one cannot establish the function of every editing site, and, in order for the preadaptation explanation to be applicable here, we have to extend the meaning of "function" in the definition of preadaptation to a broader term, e.g. "phenotypic manifestation" of a nucleotide, which would include traits such as the occurrence of a specific amino acid at a specific position of a protein.

#### 3. Caveats

**3.1. Underpredicted editing sites.** The procedure to identify editing sites employed by Liscovitch-Brauer et al. is based on the alignment of RNA and genomic reads to constructed transcripts and the analysis of mismatches (Liscovitch-Brauer et al. 2017). Editing sites in transcripts with low read coverage are likely to be missed, as the average editing level does not exceed 10% in all studied species. As we calculate the substitution matrices using substitutions that happened between the ancestral and descendant states, the former inferred from at least two species, this could result in underprediction of ancestral editing sites, which, in turn, would inflate the number of A-to-E transitions. However, for NES we observe the same behavior as for the majority of sites (Fig. 3), but for SES the effect is much weaker, even considering the fact that they are generally less conserved than NES, while if the underprediction had determined our results, we would observe the same picture for NES and SES.

**3.2. Heterozygous editing sites**. During construction of the editing-site list, heterozygous sites A-G were not considered (Liscovitch-Brauer et al. 2017), and hence some editing sites could not be predicted. However, heterozygous sites influenced the constructed transcriptomes, which by definition contained the allele prevailing in the reads (Liscovitch-Brauer et al. 2017). This could influence the results in two ways.

Firstly, consider a heterozygous A-G site in one species and a homozygous G-G site in another species. Clearly, there has been a G-to-A transition; at that, the procedures for the construction of the transcriptome and edited site list would generate the former transcript strictly with either G or unedited A. In the former case the substitution would not be counted, and in the latter case we would count a G-to-(unedited)A transition. If the adenine in the heterozygous state is edited, the above procedure would report either no transition of this adenine, or a transition involving an unedited adenine, hence decreasing the real number of E-to-G. Similarly, in the case of A-G and A-A sites, the G-to-E substitutions would also be undercounted.

Secondly, the lack of data about heterozygous editing sites would result in a general underprediction of A-to-I editing sites discussed above. Thus, this underprediction of E-to-G and G-to-E transitions could influence the calculated $R_G$ values, but it would act against the reported effects, as it lowers the $p(E \rightarrow \mathrm{G})$ values.

#### 4. Supplementary Figures

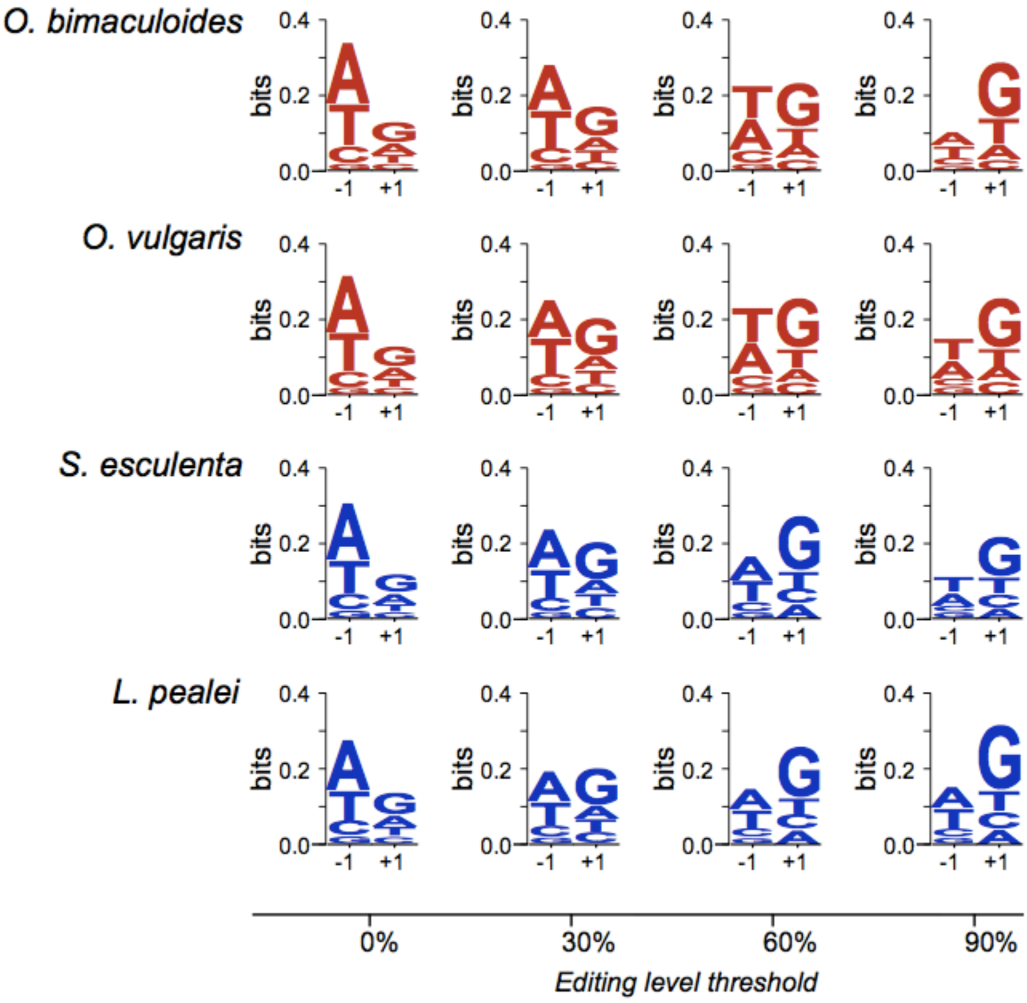


**Supplementary Figure S1 | LOGOs of nucleotides adjacent to editing sites in all studied organisms at different editing levels.**

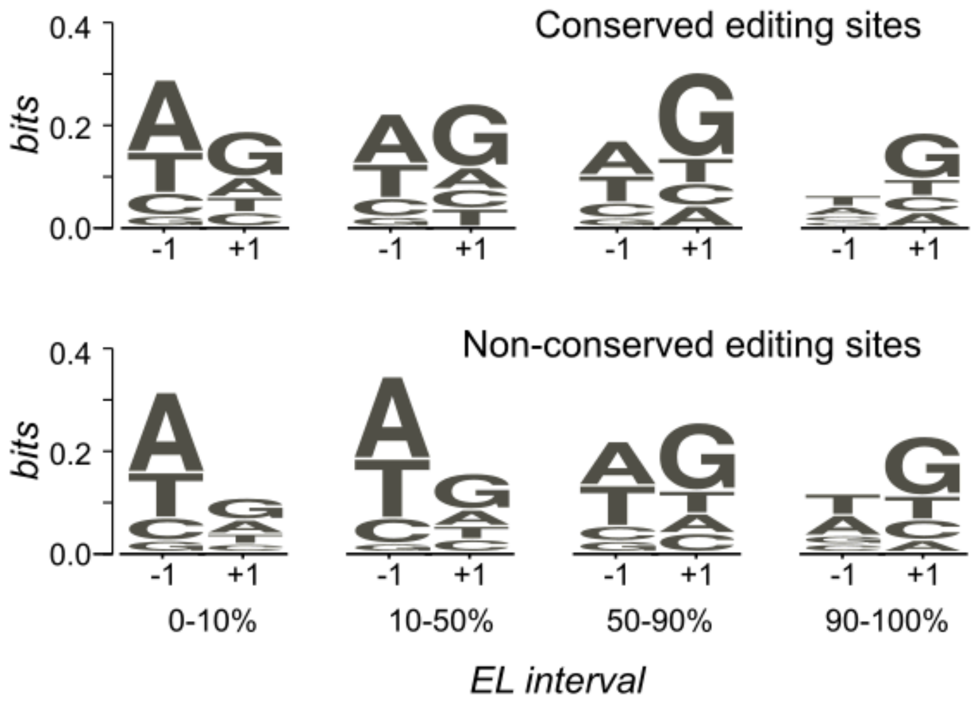


**Supplementary Figure S2 | LOGOs of nucleotides adjacent to conserved and non-conserved editing sites in the squid-cuttlefish pair.**

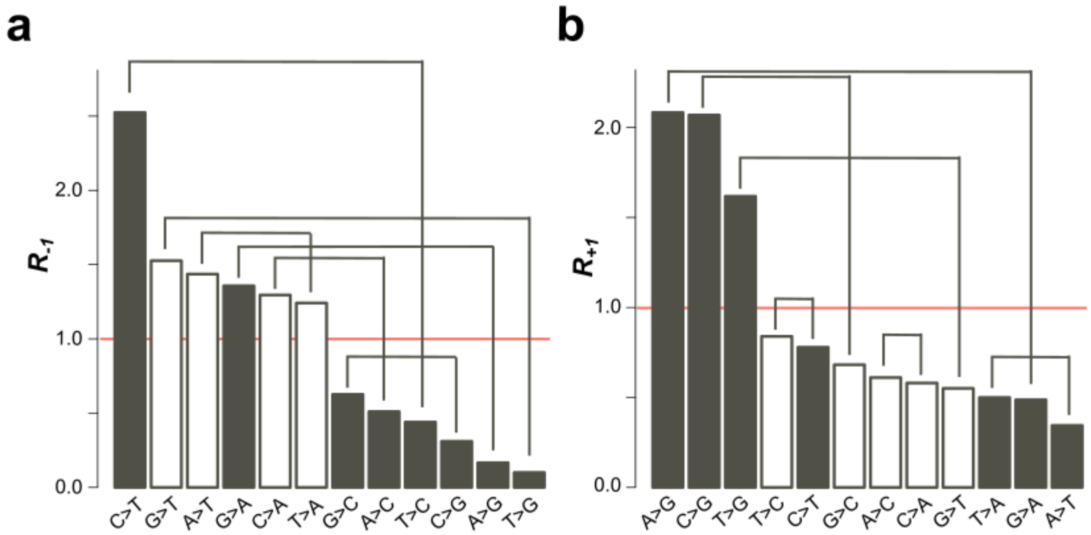


**Supplementary Figure S3 | Over- and underrepresented mismatch *R* values in the local context of non-conserved editing sites in the pair of *Octopus* species.** *R* values are defined for the ±1 positions relative to transcriptomic adenines as the ratio of the probability of a given substitution near the edited adenine and the respecteve probability for the non-edited adenine. **(a)** In position –1 relative to NCES sites. **(b)** In position +1 relative to NCES sites. Lines represent mutually reversed substitutions. White bars represent *R* values which do not significantly differ from 1 ($p > 0.05$) and grey bars represent statistically significant *R* values.

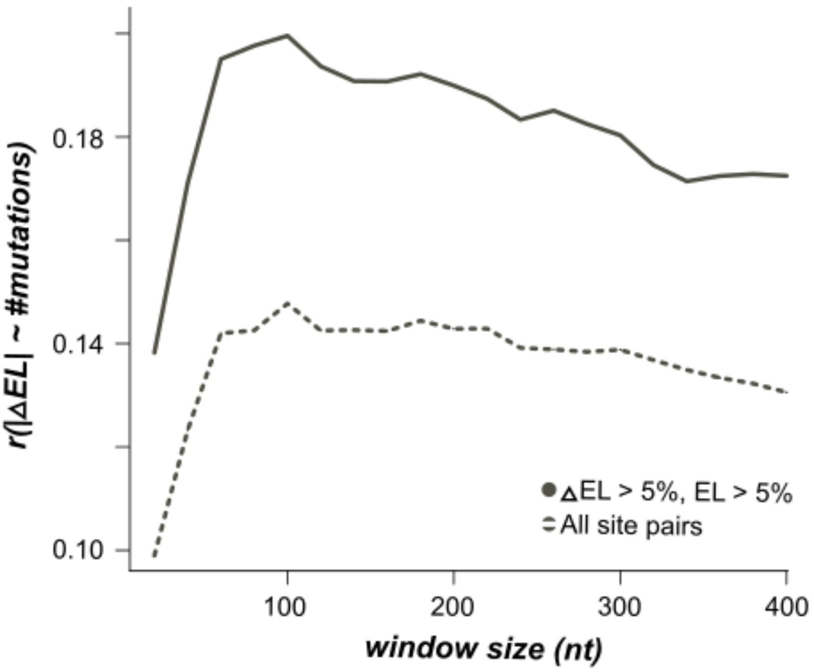


**Supplementary Figure S4 | Pearson's correlation between the absolute value of difference of ELs in homologous sites and the number of mismatches in a window of a given size for different window sizes.** The dashed line represents values obtained for all homologous site pairs; the solid line represents values obtained for pairs of homologous edited adenines with EL above 5% and with the absolute value of EL difference greater than 5%.

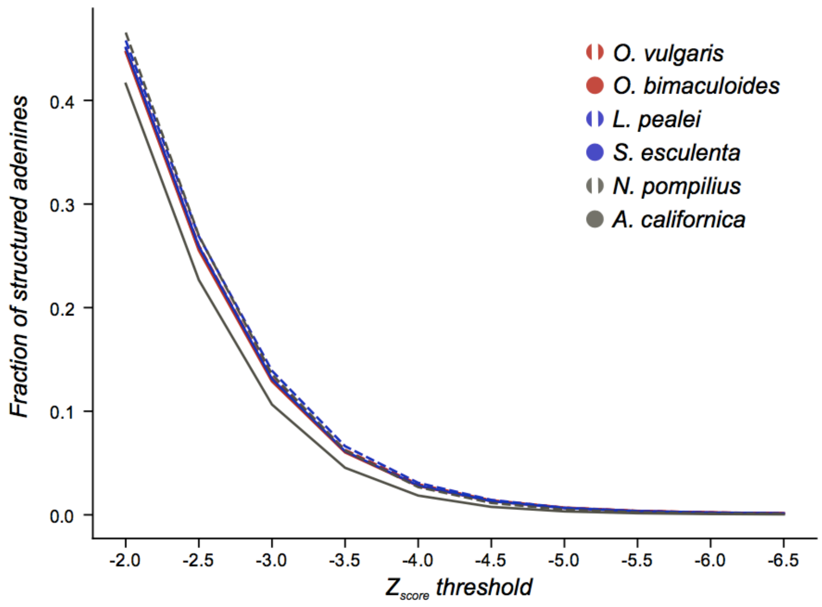


**Supplementary Figure S5 | *A. californica* has less adenines in structured regions then other species.** The classification of structured and unstructured segments was performed with various thresholds (see Methods). The grey line corresponding to *A. californica* is considerably lower than cephalopod lines.

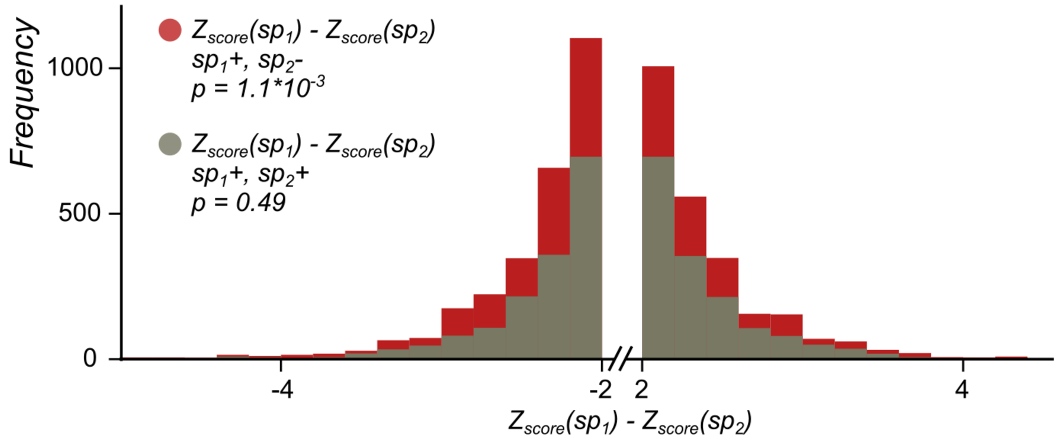


**Supplementary Figure S6 | The local secondary structure is more stable at edited adenines than at homologous, non-edited adenines.** The distribution of the difference of the minimal free energy Z-score between homologous sites in two octopuses, *O. vulgaris* and *O. bimaculoides* is shown in red when two homologous sites have different editing status (edited minus unedited) and in grey when both sites in a pair are edited. The left tail of the red histogram is heavier than the right one ($p$ = 6.03×10$^{-17}$ versus 0.57 for the grey histogram), showing that the editing sites tend to regions with higher secondary structure stability.

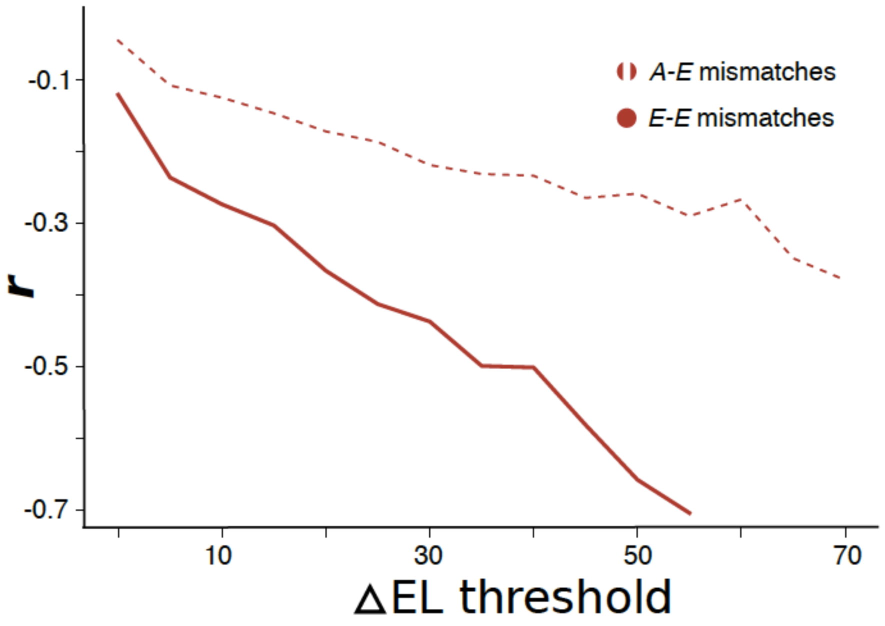


**Supplementary Figure S7 | Dependence of Pearson's correlation between the difference of structural Z-scores and the difference in ELs of homologous sites on the minimal considered difference in ELs.** The dashed line represents values obtained for pairs on homologous adenines, such that one adenine in a pair is edited, and the other is not. The EL of all unedited adenines was set to 0. The solid line represents values obtained for pairs of homologous edited adenines. Only correlation coefficients with p-values below 0.05 are shown.

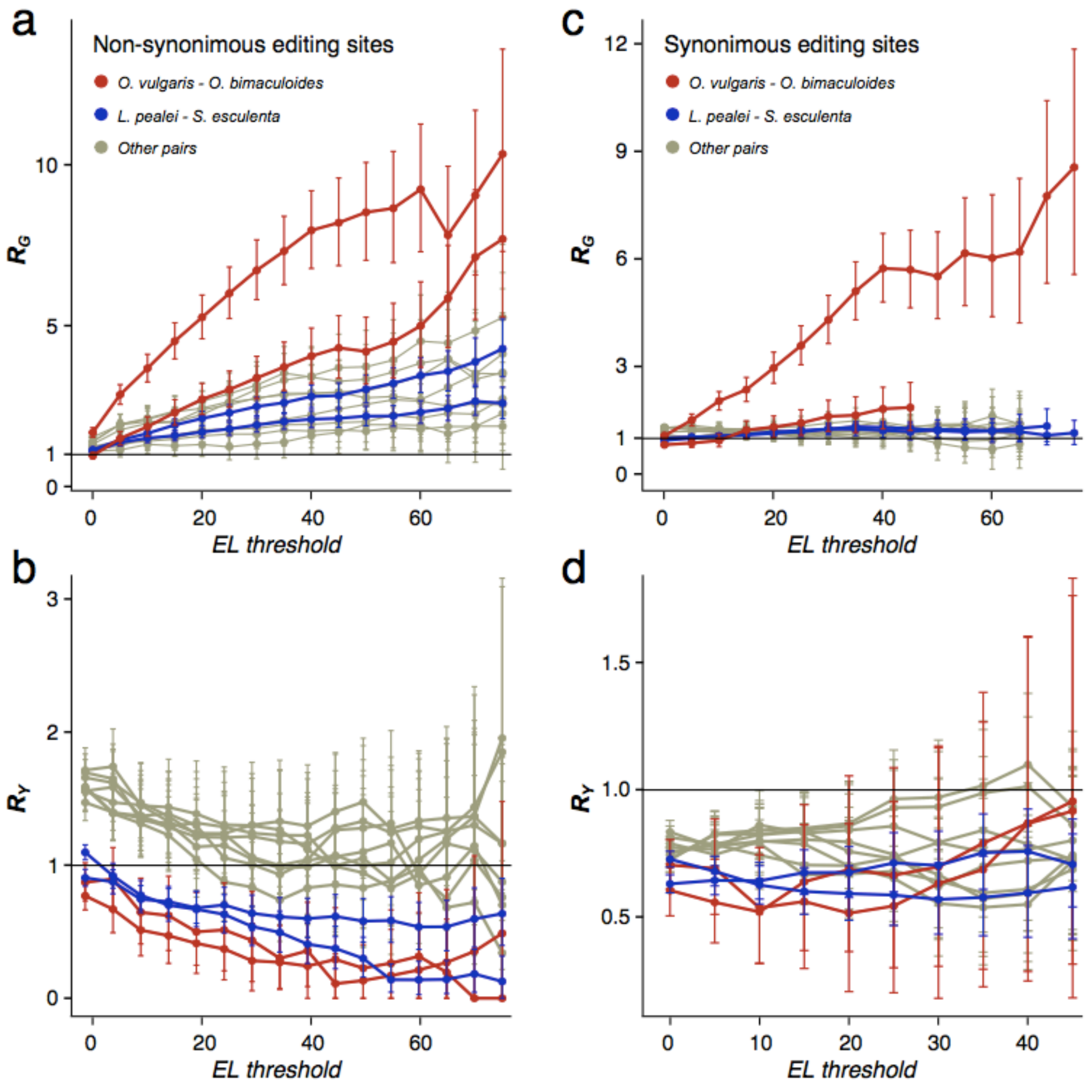


**Supplementary Figure S8 | Dependence of $R_G$ and $R_Y$ on the editing level considered separately for non-synonymous (ab) and synonymous (cd) sites.** Two curves for each pair are given, since $R_N$ is a reference-based measure. The red curves correspond to the *Octopus* pair, the blue curves, to the pair cuttlefish–squid, the grey curves, to distant pairs.

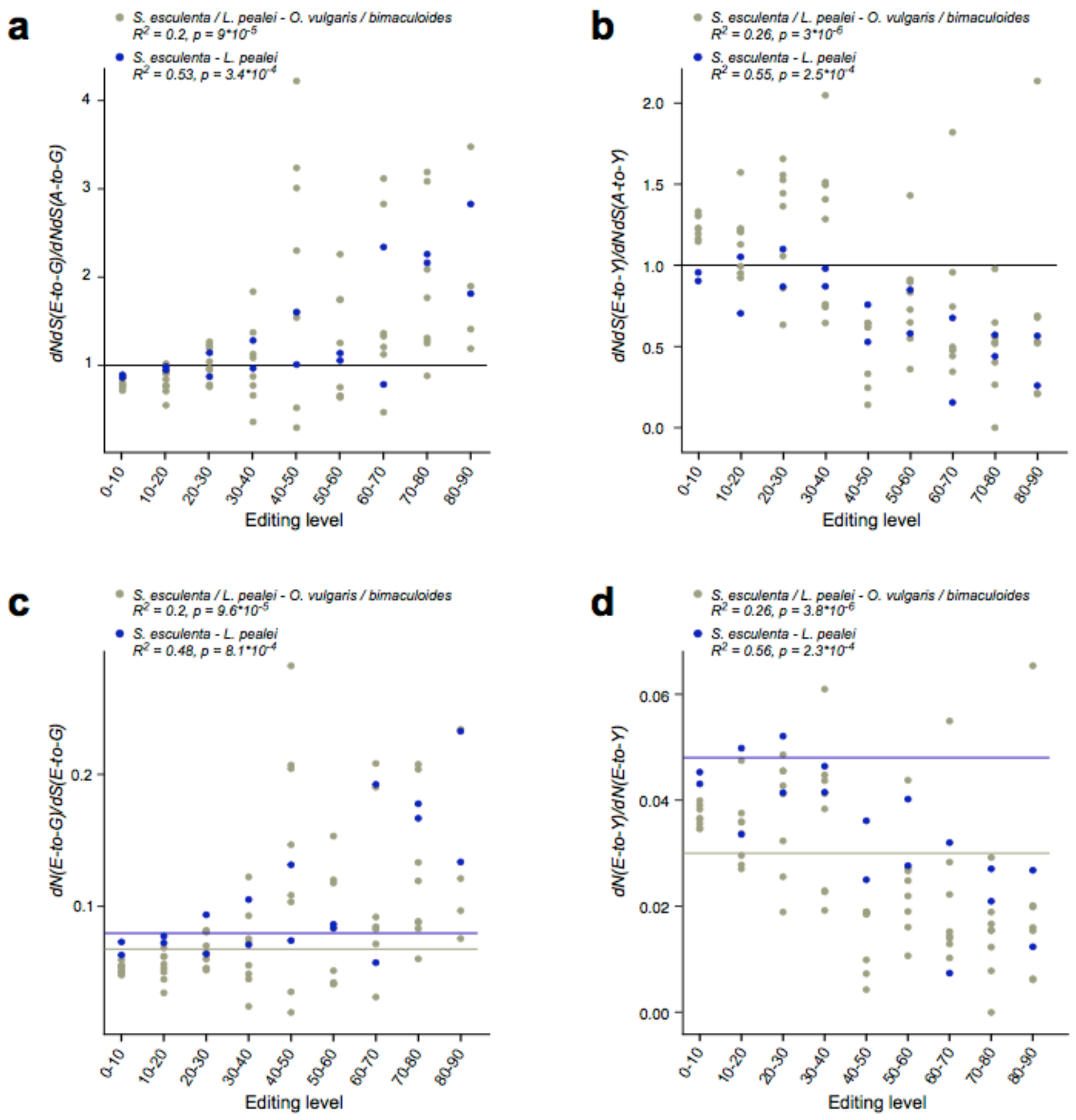


**Supplementary Figure S9 | Normalized *dN/dS* of editing site substitutions**. *dN/dS* of editing site substitutions normalized by respective *dN/dS* of unedited adenines to G **(a)** and to pyrimidines **(b)**. *dN/dS* of editing sites normalized by the respective $\xi^{non}/\xi^{syn}$ value ratio (Supplementary Table 1) for substitutions to G **(c)** and to pyrimidines **(d)**. Horizontal lines represent average *dN/dS* values for unedited adenines. $R^2$ values and *p*-values calculated from the *F* statistic are shown.

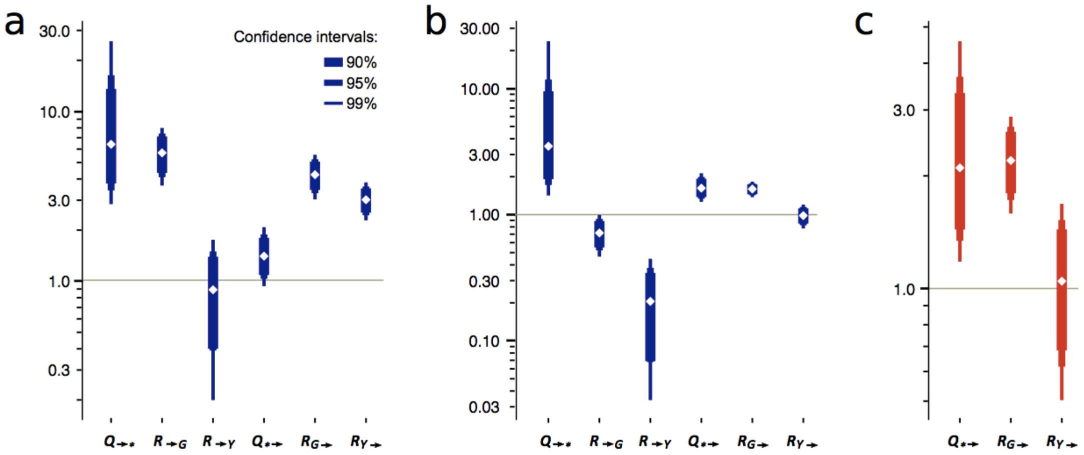


**Supplementary Figure S10 | Mutational characteristics of editing sites. (a,b)** Mutational characteristics of edited sites for the squid–cuttlefish pair separately for non-synonymous (NES) **(a)** and synonymous (SES) **(b)** sites. **(c)** $Q_{*\rightarrow}$ and $R_{*\rightarrow}$ values calculated for the octopus total substitution matrix; there are insufficient data for the separate analysis of NES and SES.
